# Protein design to broadly reprogram engineered T cell function

**DOI:** 10.64898/2026.08.13.742806

**Authors:** Scott E. Boyken, Sean Merillat, Robert A. Langan, Howell F. Moffett, Brian Coventry, Françoise Haeseleer, Kevin G. Haworth, Inna Goreshnik, Joseph DeSautelle, John Chukinas, Bradley Hammerson, Thaddeus M. Davenport, Duy Nguyen, Rupesh Amin, Shujun Yuan, Glenna Wink Foight, Brian D. Weitzner, Aaron E. Foster, David Baker, Marc J. Lajoie

## Abstract

The efficacy of engineered T cell therapies in solid tumors remains limited by T cell dysfunction, driven by complex processes that cannot be easily manipulated via genetic knockouts or overexpression of individual genes. Protein design can create new biological functions that can rewire these consequential cell fate decisions. Here, we introduce OUTLAST Regulators, designed proteins that reprogram critical T cell signaling pathways to enhance functional persistence. These proteins are capable of regulating diverse groups of proteins such as the NR4A family of pro-exhaustion transcription factors, E3 ligases Cbl-b and c-Cbl, and SOCS family proteins. Our designs markedly improve CAR-T and TCR-T performance *in vitro* and *in vivo* in stringent solid tumor preclinical models. OUTLAST Regulators are implemented as compact genetic modules compatible with standard viral vectors and cell therapy manufacturing processes, creating a powerful platform for programming new functions into enhanced cell and gene therapies.

## Introduction

Cell therapies hold immense promise to treat intractable diseases by sensing and responding to complex biology in ways that conventional small molecule and protein-based drugs cannot. Current approaches to enhance cell therapy function are limited in their ability to harness this multifaceted potential because they have largely relied on modifying single endogenous genes, and attempts to overexpress or knock out multiple genes face translational hurdles around vector payload capacity and multi-allele editing efficiency. Engineered immune cell therapies exemplify both the promise and challenge of these approaches: chimeric antigen receptor (CAR) T cells revolutionized the treatment of hematologic malignancies^1^, but efficacy in solid tumors has been limited by poor functional persistence due to a hostile tumor microenvironment and the rapid onset of T cell exhaustion^2^. Solid tumors drive T cell dysfunction redundantly through many independent mechanisms that cannot be overcome by disrupting a single target protein; small molecule regulation^3^, CRISPR knockouts (KO)^4–6^, static gene overexpression^7^, and bioPROTACs^8,9^ to degrade individual targets^10^ have yielded improved CAR-T cell functions but fall short of adequately addressing these myriad challenges. We hypothesized that this problem could be solved by designing novel protein regulators that more comprehensively regulate T cell function by modulating multiple signaling proteins simultaneously.

Here we describe the use of *de novo* protein design to create OUTLAST Regulators, compact modules that interact with multiple intracellular targets to reprogram cell function, with the goal of improving functional persistence, preserving stemness and reducing T cell exhaustion (**Figure 1**). To maximize therapeutic potential, we set out to achieve designs that are small enough to be modularly encoded in standard viral vectors (<1kb DNA), compatible with conventional cell therapy manufacturing processes, and capable of enhancing both chimeric antigen receptor (CAR) and transgenic T cell receptor (TCR) T cell function across different solid tumor models. To select target families of interest, we started with proteins known to be associated with T cell dysfunction and for which gene knockouts improved CAR T cell cytotoxicity and cytokine production after repeated stimulation in a tumor co-culture assay (**Figure S1**). Based on this evaluation, we chose to focus on the nuclear receptor subfamily 4A (NR4A) pro-exhaustion transcription factors^11^, the Casitas B-lineage lymphoma (Cbl) ubiquitin ligases Cbl-b and c-Cbl^12^, and the Suppressor of Cytokine Signaling (SOCS) family^13^.

**Figure 1.**
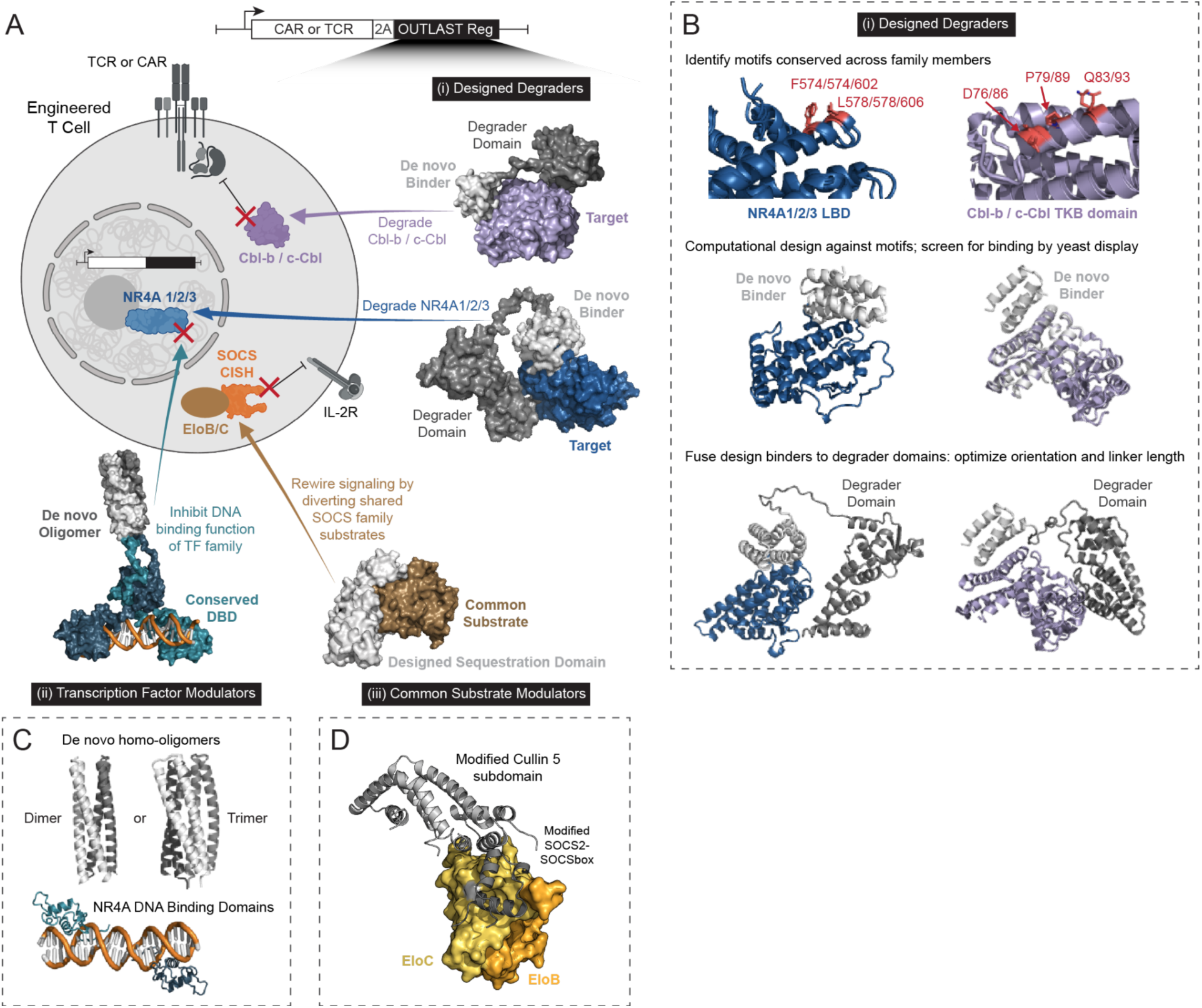
OUTLAST Regulator design strategies to broadly reprogram T cell function. (A) OUTLAST Regulators are compact modules that can be genetically encoded into engineered cells via standard vectors: (i) Designed Degraders are heterobifunctional proteins consisting of a *de novo* binder and an engineered degrader domain; (ii) Transcription Factor (TF) Modulators oligomerize conserved DNA-binding domain (DBD) motifs to outcompete endogenous TFs; and (iii) Common Substrate Modulator rewire signaling pathways by diverting shared protein substrates. T cell illustration was created using BioRender (https://biorender.com, Fontana, T. 2024); all structural images rendered using PyMOL. (B) Workflow to create Designed Degraders targeting NR4A1/2/3 and Cbl-b / c-Cbl. Conserved motif residues shown in red for NR4A1/2/3 LBD (blue, models based on PDB 3V3E) and Cbl-b / c-Cbl (purple, models based on PDB: 3VGO, Cbl-b residues 39-426); computational design generated *de novo* binders against conserved motifs; designs were screened for binding by yeast display; binders were subsequently fused to E3 ligase degrader domains (Figure S2), and orientation and linker lengths were optimized. (C) pan-NR4A Transcription Factor (TF) Modulators were designed by combining *de novo* homodimers or trimers to the NR4A1 DNA Binding Domain (DBD) with various orientations and linker lengths. (D) SOCS family Common Substrate Modulator was designed by fusing a modified Cullin5 subdomain to a modified SOCS2-SOCSbox.

We created three distinct classes of OUTLAST Regulators (**Figure 1A**). First, Designed Degraders are heterobifunctional proteins where binders to target family epitopes are fused to engineered degrader domains in a manner that induces proteasomal degradation of target family members upon binding; we used computational protein design to create *do novo* binders against conserved motifs that enable multiple target proteins to be regulated simultaneously by a single compact Designed Degrader, and we applied this approach to create designs that regulate the entire family of NR4A1/2/3 and Cbl-b/c-Cbl (**Figure 1B**). Second, Transcription Factor (TF) Modulators are designed proteins that leverage *de novo* oligomers to pre-organize conserved DNA-binding motifs and inhibit the function of endogenous TFs, and we applied this approach to create designs that regulate NR4A1/2/3 (**Figure 1C**). Third, Common Substrate Modulators are redesigned protein interaction domains that rewire signaling pathways by inhibiting or diverting shared signaling substrates, and we applied this approach to create designs that regulate the SOCS family (**Figure 1D**). The TF Modulator and Common Substrate Modulator regulation strategies allow for control of functions beyond degradation, and we demonstrate that each of these three design strategies can substantially enhance engineered T cell function in solid tumor models.

## Results

### Design and characterization of pan-NR4A OUTLAST Regulators

The NR4A family of transcription factors consists of NR4A1 (NUR77), NR4A2 (NURR1), and NR4A3 (NOR1), all of which have been shown to play key roles in driving T cell dysfunction by suppressing effector function, activating co-inhibitory receptors, and inducing exhaustion^11^. In studies of murine CAR T cells, a triple KO of all three family members enhanced antitumor function^4^. The triple KO exceeded the benefit of any individual KO, with NR4A3 KO showing the largest effect of any individual KO^4^, a result that was recapitulated in NR4A KO studies in human T cells^14^. A recent approach that combined a CRISPR gene knock-in with a small molecule PROTAC showed that the ability to regulate NR4A family members can reduce exhaustion and improve anticancer T cell function^15^. Despite showing functional promise, these gene KO and knock-in approaches require editing several alleles at once, posing challenges for cell therapy manufacturing and translatability.

We set out to create OUTLAST Regulators capable of simultaneously down-regulating the entire family of NR4A TFs without requiring cumbersome gene editing. We started with the Designed Degrader approach and computationally designed protein binders for a structural motif at the C-terminus of the ligand-binding domain (LBD) that we identified to be conserved across all NR4A family members (**Figure 1B**). Protein backbones were generated around the LBD motif using RFDiffusion^16^, amino acid sidechains designed with ProteinMPNN^17^, and computational designs filtered and ranked using analysis metrics from Rosetta^18^ and AlphaFold2^19^ predictions (**Figure S3A; see Methods**). 7,500 designs were tested by yeast surface display to assess binding against purified recombinant LBD of NR4A1 and NR4A3; cells were sorted by FACS to identify binder hits against NR4A1 and NR4A3 individually, and then these hits were sorted by FACS to identify designs that bound both targets (**see Methods**). Hundreds of designs showed high-affinity binding to both NR4A1 and NR4A3; these designed binders are named “nr##” and we selected 24 binder designs estimated to have low nM K_d_ affinity for further characterization (**Figure 2A, Figure S3A**). In parallel, we screened a panel of E3 ligase domains for the ability to efficiently degrade NR4A proteins in primary T cells. To facilitate rapid screening and assess degrader domain function in the absence of binders, we developed an assay that leveraged *de novo* designed heterodimers (DHDs)^20^ to directly bring overexpressed HA-tagged NR4A target protein into close proximity with different degrader domains (**Figure S2**). FBXL12, RNF8, SOCS2_143-198_ (“SOCS2box”), and SPOP_168-364_ (hereafter referred to as “SPOP”) all showed promising activity against NR4A targets in this screen (**Figure S2**).

**Figure 2.**
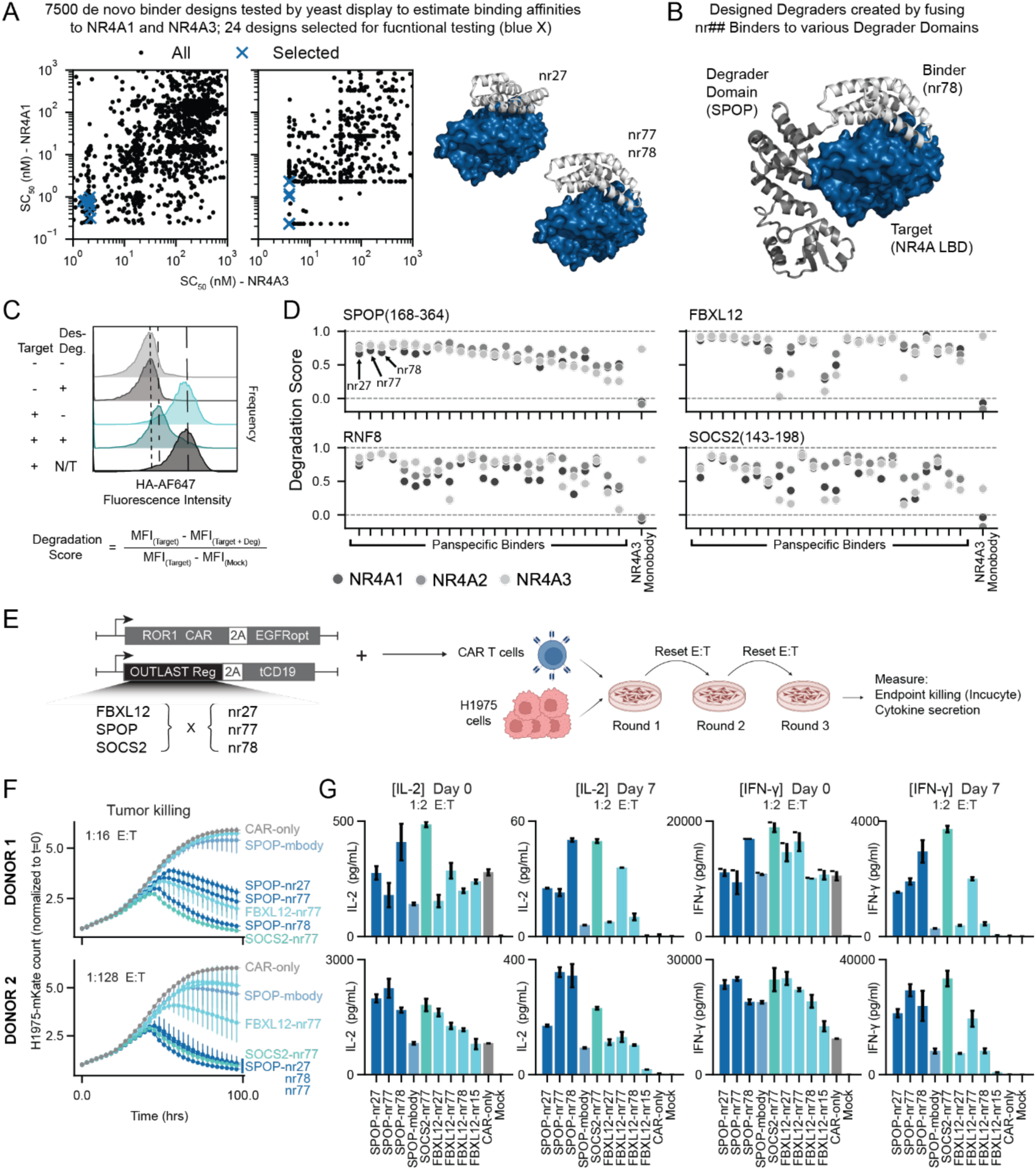
***In vitro* function of pan-NR4A Designed Degraders.** (A) Estimated binding affinities from yeast display: lower-bound (*left*) and upper-bound (*right*) of the midpoint concentration (SC_50_) in the binding transitions for each design; 24 designs selected for functional characterization are indicated by blue crosses. Structural models of representative binder designs nr27, nr77, and nr78 shown in cartoon representation (PyMOL) bound to NR4A1 LBD (blue); nr77 and nr78 share an identical backbone but have distinct amino acid sequences. (B) Representative Designed Degrader SPOP-nr78: AlphaFold2^19^ structural model of nr78 fused to the C-terminus of SPOP_168–364_ showing predicted binding geometry between nr78, SPOP, and NR4A LBD. **(C-D)** Primary human T cells were dual-transduced with a lentiviral vector expressing the designed Designed Degraders and a second vector expressing HA-tagged NR4A1, NR4A2, or NR4A3 target; T cells transduced with individual vectors served as controls. Cells were stained by FACS monitoring HA-AF647 to assess target degradation in the presence or absence of the Designed Degraders. (C) Representative flow cytometry data quantifying HA-tagged NR4A levels in the presence or absence of the Designed Degrader (“Des-Deg”). “Degradation Score” is derived by mean fluorescence intensity (MFI) values and the formula shown; a Degradation Score of 1.0 indicates complete degradation. (D) Degradation Score for Designed Degrader designs against NR4A1, NR4A2, and NR4A3; 24 selected binders from (A) were fused to four different degrader domains: SPOP_168–364_ (“SPOP”), FBXL12, RNF8, and SOCS2_143-198_ (“SOCS2box”). The NR4A3-specific Designed Degrader (“SPOP-Monobody”) is included as a control. (E) Schematic of serial tumor restimulation assay used to evaluate the functional activity of CAR-T cells engineered with pan-NR4A Designed Degrader designs; one vector encodes a ROR1-targeted CAR with a transduction marker (EGFRopt^40^), and the second vector encodes the designs with a tCD19 transduction marker. Engineered CAR-T cells were subjected to multiple rounds of antigen-specific stimulation on H1975 tumor cells, followed by endpoint tumor killing and cytokine detection assays. (F) Incucyte-based quantification of H1975 tumor cell killing kinetics following 3 rounds restimulation. (G) IL-2 and IFN-γ cytokine secretion measured at Day 0 and Day 7 after three rounds of restimulation

Designed Degraders were constructed from the 24 chosen binders fused to each of the four ligase domains (**Figure 2B**). Lentiviral vectors encoding the Designed Degraders with a truncated CD19 (tCD19) transduction marker were dual-transduced into primary human T cells together with a second lentiviral vector expressing HA-tagged NR4A1, NR4A2, or NR4A3 target with an EGFR transduction marker. T cells transduced with only one of the vectors served as controls. After 3 days of cell expansion, cells were stained intracellularly for the HA-tag along with surface staining of tCD19 and EGFR, and a “Degradation Score” was calculated by measuring the mean fluorescence intensity (MFI) of the Target in the presence or absence of the Designed Degrader, normalized to a background control (**Figure 2C**). This metric is used throughout this study to compare the relative activity of different Designed Degraders, and a Degradation Score of 1.0 indicates complete degradation of detectable target protein. All 24 designed binders tested were able to degrade all three NR4A family members (Degradation Score >0.5) when paired with at least one of the chosen degrader domains (**Figure 2D**). For the Designed Degraders with SPOP or FBXL12, ∼80% (19/24) had degradation scores >0.5 for all family members, and 75% (18/24) of the FBXL12 designs had Degradation Scores >0.75 for all family members, with many designs approaching 1.0 (**Figure 2D**). These success rates are remarkable because the binders were functional straight from the computer and did not require any further optimization based experimental results.

We next set out to evaluate the ability of these designs to enhance functional persistence by degrading endogenous NR4A family members in the context of CAR T cells targeting receptor tyrosine kinase-like orphan receptor 1 (ROR1), an antigen relevant for numerous solid tumor cancers^21^. Primary T cells were dual-transduced with lentiviral vectors encoding the Designed Degrader designs along with a lentiviral vector encoding a 2nd-generation ROR1 CAR (R12-CD28TM-41BB-CD3z)^22^ and assessed for tumor killing and cytokine production after three rounds of serial restimulation by tumor co-culture against ROR1^+^ H1975 cells (**Figure 2E**). CAR T cells become dysfunctional in response to chronic antigen stimulation over time, losing their ability to kill tumor cells and produce cytokines^23^, and we leverage this repeated tumor challenge assay throughout to identify designs with enhanced *in vitro* functional persistence. The stringency of this assay can be increased by reducing the effector to target (E:T) ratio, *i.e.* the number of effector CAR T cells relative to tumor cells (**Figure 2F**), and by increasing the rounds of tumor challenge. T cells expressing ROR1 CAR alone were able to control tumor cells in early rounds of coculture but failed after multiple rounds at lower E:T; at the start of the experiment, ROR1 CAR T cells produced similar levels of IFN-γ and IL-2 compared to CAR T cells with co-expression of the Designed Degraders, but by the end of the challenge were unable to produce detectable levels of these cytokines (**Figure 2G**). The Designed Degrader designs all enhanced tumor cell killing and cytokine production (**Figure 2F-G**), consistent with improved functional persistence. To assess the functional impact of regulating NR4A3 alone vs all three family members, we generated a monobody that specifically binds to NR4A3 with an estimated K_d_ of ∼9 nM but that does not bind to NR4A1 or NR4A2 (**Figure S4**). Designed Degraders that incorporated this monobody specifically degraded NR4A3 but not NR4A1 or NR4A2 (**Figure 2D**). Functionally, NR4A3-specific Designed Degraders improved ROR1 CAR T cell cytotoxicity and cytokine production after chronic stimulation compared to the ROR1 CAR alone but did not perform as well as the pan-NR4A Designed Degraders (**Figure 2F-G**). SPOP-nr78 and SPOP-nr77 designs performed the best in these initial experiments and were chosen for deeper characterization.

Biolayer interferometry (BLI) confirmed that nr78 and nr77 binder designs bound the LBD of all three NR4A family members with an estimated K_d_ of ∼1 nM (**Figure S3B-C**), a striking result given that no affinity maturation was required. To verify the computational design models in the absence of experimentally determined co-complex structures, we mutated binding interface residues to amino acid identities expected to disrupt target binding without affecting soluble expression or folding; as expected, these mutations ablated binding (**Figure S3B-C**) and resulted in no degradation of NR4A target protein in T cells (**Figure S3D**). To verify that degradation was occurring in the T cells via ubiquitin-dependent proteasomal pathways expected for the SPOP (Speckle-type POZ protein)^24^ degrader domain, which is a substrate-binding adaptor for the Cullin RING ligase 3 (CRL3) complex^25^, we monitored degradation in the presence of a compound known to block Cullin-mediated proteasomal activity, the pan-neddylation inhibitor MLN4924^26^. As expected, the addition of MLN4924 prevented target degradation by Designed Degrader designs that contained SPOP (CRL3), FBLX12 (CRL1), and SOCS2box (CRL5), whereas RNF8 (a RING-finger domain E3 ubiquitin ligase that is not mediated by Cullin-RING ligases) was unaffected (**Figure S5**).

As an orthogonal mechanism to regulate NR4A family function, we designed TF Modulators that employ *de novo* designed protein homo-oligomers to regulate NR4A function at the DNA-binding level. The DNA binding domains (DBDs) of the NR4A family are highly conserved and exert function through binding known DNA response elements as monomers and cooperative homo- or hetero-dimers^27^. We hypothesized that pre-organized oligomers of NR4A DBDs (lacking the LBD and transactivation domain) could outcompete endogenous NR4A proteins at these DNA binding sites, thus inhibiting NR4A function. We generated designs using *de novo* homodimer and homotrimer helical bundles^28,29^ fused to the DBD of NR4A1 (**Figure 1C, Figure S6A**). All of the TF Modulator designs improved CAR T cell cytotoxicity and cytokine production compared to the ROR1 CAR alone (**Figure S6B-D**). The dimerized DBD performed comparable to a tandem repeat of the DBD included as a control (**Figure S6B**), and the trimerized designs improved tumor killing and cytokine production compared to the tandem repeat design (**Figure S6C**). The strength of the functional effect was dependent on multiple design features such as linker length, fusion orientation, and domain cutoff of the DBD; for example, fusing the homo-trimer to the N-terminus of the DBD outperformed fusing it to the C-terminus (**Figure S6C**), and an extended version of the DBD (NR4A1 residues 263-399) outperformed a shorter version (NR4A1 residues 263-358) (**Figure S6D**). To confirm the designed mechanism, we mutated key DNA-binding residues on the TF Modulators and found that this mutant abrogated all CAR T cell functional benefits (**Figure S6D**). Of the TF Modulators tested, the best design connected a homotrimerizing domain to the N-terminus of the extended NR4A DBD (NR4A1 residues 234-399) via a 5xG4S linker, and this “Trimer-NR4A-DBD” design was chosen for further functional characterization (**Figure 3**).

**Figure 3.**
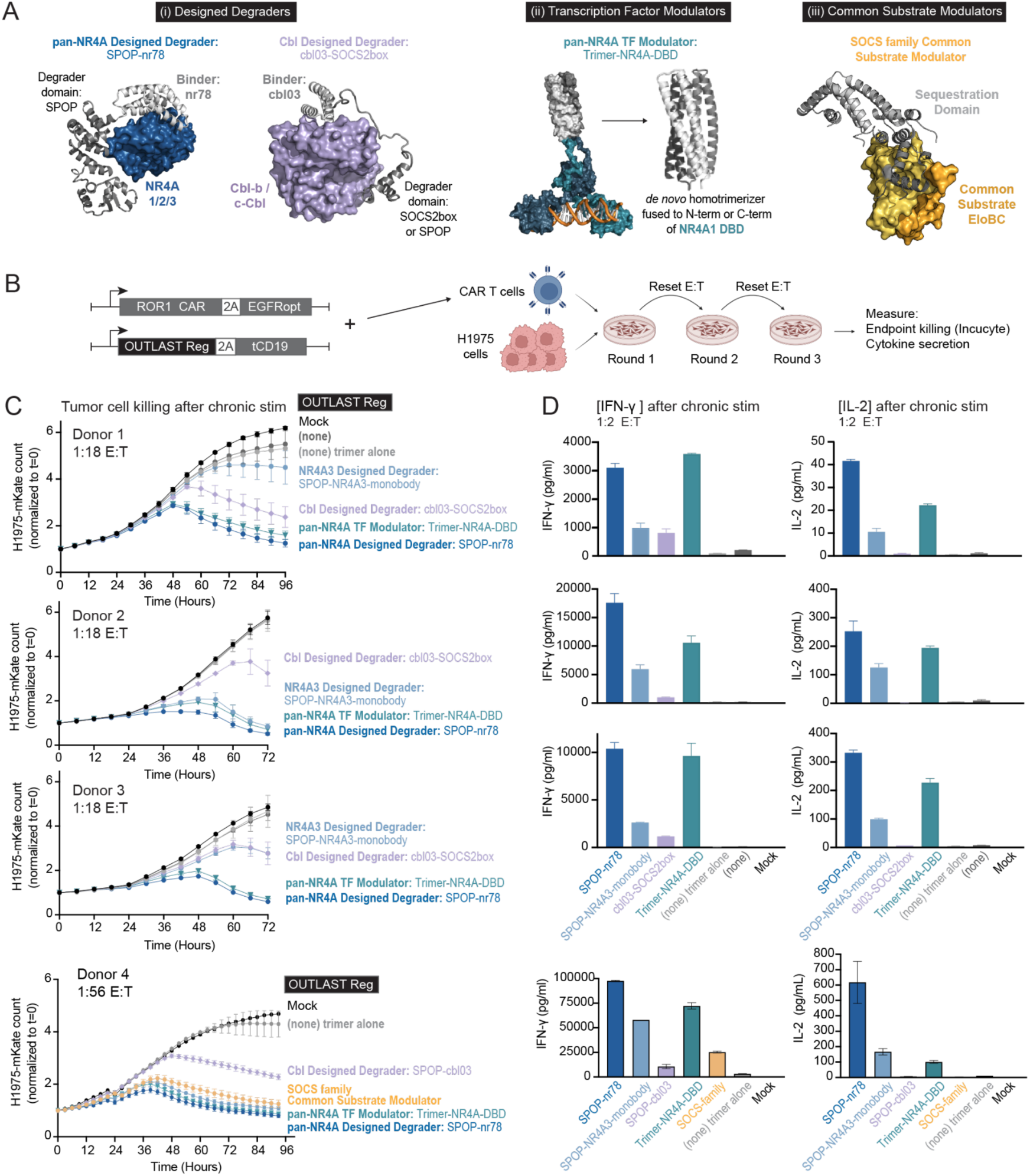
***In vitro* CAR-T function comparing best designs from each OUTLAST Regulator class.** (A) Structural models for the best performing designs from each class: pan-NR4A Designed-Degrader (Figure 2), Cbl Designed Degrader (Figure S7), pan-NR4A TF Modulator (Figure S6), and SOCS family Common Substrate Modulator (Figure S8). (B) OUTLAST Regulator designs were dual-transduced in T cells along with the ROR1 CAR and CAR-T cell function was evaluated by serial restimulation in a tumor co-culture assay; experimental diagram created using BioRender (https://biorender.com, Fontana, T. 2024). Control constructs: “none” = ROR1 CAR-T cells without an OUTLAST Regulator on the second plasmid; “trimer alone” = CAR-T cells transduced with an inert designed trimer on the second plasmid; “Mock” are non-transduced T cells. (C) Incucyte-based quantification of H1975 tumor cell killing kinetics after three rounds tumor challenge (D) IFN-γ and IL-2 cytokine secretion measured at Day 7 after three rounds of tumor challenge.

### Design and characterization of Cbl OUTLAST Regulators

Cbl-b and c-Cbl are RING-type E3 ubiquitin ligases that negatively regulate T cell activation, maintaining immune tolerance by promoting the degradation of proximal signaling molecules like CD3ζ, ZAP-70, and Vav1^30^. Cbl-b KO CAR T cells have been shown to improve CAR T cell function^31^. We aimed to generate Designed Degraders that regulate Cbl-b and c-Cbl using the same general design and screening strategy that we applied to create the pan-NR4A Designed Degrader designs. We identified a motif that is conserved across the Cbl-b and c-Cbl N-terminal Tyrosine Kinase Binding (TKB) domain, but distinct from other related domains, and we computationally designed *de novo* binders against these motifs (**Figure 1B**); this motif is also conserved in the homologue Cbl-c, but because Cbl-c is generally not expressed in T cells^32^, we did not include it in testing for this study. Using yeast display, we selected binder designs (named “cbl##”) with low-nanomolar affinity to Cbl-b and measured binding kinetics by BLI for the three designs with the strongest affinity (**Figure S7A**). We then fused these three binders to degrader domains capable of down-regulating Cbl-b (**Figure S2**) and evaluated their ability to degrade Cbl-b target protein in primary T cells. Target degradation was assessed as described above, and multiple Cbl Designed Degraders were shown to robustly degrade Cbl-b in T cells (**Figure S7B-C**). Binder cbl03 had the strongest affinity with an estimated K_d_ of 0.22 nM (**Figure S7A**), and when cbl03 was fused to SOCS2_143-198_ (“SOCS2box”) or SPOP degrader domains, it degraded both Cbl-b and c-Cbl targets in T cells (**Figure S7D**). To verify the cbl03 computational design model, we mutated interface residues, which ablated the degradation activity for both Cbl-b and c-Cbl as expected (**Figure S7D**). Cbl Designed Degraders with the cbl03 binder fused to SOCS2box or SPOP were chosen for deeper functional characterization and improved ROR1 CAR T cell cytotoxicity and IFN-γ production after chronic antigen stimulation when co-expressed compared to ROR1 CAR alone (**Figure 3B-D**).

### Design and characterization of SOCS family OUTLAST Regulators

SOCS family proteins, which include SOCS1-7 and CISH (Cytokine-Inducible SH2-containing protein), are negative regulators of cytokine receptor signaling (via SOCS1-3 and CISH) and receptor tyrosine kinase (RTK) signaling (via SOCS4-7), that inhibit signaling through binding of their SH2 domains or by acting as a substrate recognition subunit for a Cullin5-based E3 ubiquitin ligase complex^13^. SOCS family members carry out these functions in part via their conserved SOCS box domain binding the common substrates Elongin B and C (EloBC)^13^, and in T cells, SOCS1 and SOCS3 directly inhibit JAK kinase function^33,34^ and CISH can down-regulate PLC-γ1^35^, making them promising targets for regulation to improve T cell functional persistence^36–38^.

The SOCS family proteins possess characteristics that make them poor candidates for Designed Degraders: a lack of conserved surface-exposed hydrophobic motifs (necessary for binding) and lysine residues (necessary for ubiquitination); in screening assays, degrader domains that worked for other targets showed poor degradation activity for SOCS1 target in T cells (**Figure S2**). Thus, we focused on generating designs that modulated activity through sequestering common signaling substrates that are necessary for all SOCS family members to function. We created Common Substrate Modulator designs by creating a rigid fusion between a modified Cullin5 subdomain and a modified SOCS box domain (**Figure 1D**); the Cullin5 N-terminal domain was truncated (residues 10-192), retaining the parts that template on EloBC and SOCS box domains while ablating other protein-protein-interactions, and exposed surface hydrophobic residues were mutated to soluble amino acids; this fragment was rigidly fused to the SOCS box domain from SOCS2 such that when this design is overexpressed, it should sequester EloBC, thereby preventing the association of endogenous SOCS proteins and Cullin5 and inhibiting them from acting on their targets (**Figure S8A**). Furthermore, SOCS proteins require interaction with EloBC to stabilize intrinsically disordered regions, and inhibiting this interaction has been shown to down-regulate SOCS levels and activity^39^. By broadly inhibiting SOCS family proteins from engaging their cognate substrates, we hypothesized that we could enhance T cell functional persistence by maintaining sensitivity to JAK/STAT signaling and making the CAR T cells more responsive to lower levels of cytokines. These designs enhanced ROR1 CAR T cell activity *in vitro*, with improved tumor killing and cytokine production compared to the ROR1 CAR alone (**Figure S8B-C**). Tumor cell killing was comparable to that of the best pan-NR4A TF Modulator designs (**Figure S8B)**, although cytokine production was markedly lower than the TF Modulator (**Figure S8C**), and this design was advanced for further functional characterization (**Figure 3**).

### Multiple OUTLAST Regulator designs enhance CAR T function

We next set out to compare the various targets and types of OUTLAST Regulators head-to-head. The best designs from each class (**Figure 3A**) were evaluated for *in vitro* CAR-T function in multiple donors; for initial testing, lentiviral vectors encoding the OUTLAST Regulators were dual-transduced with lentiviral vectors encoding the ROR1 CAR, and the resulting CAR T cells were subjected to serial restimulation by repeated co-culture with ROR1^+^ H1975 tumor cells (**Figure 3B**). CAR T cells co-expressing the NR4A, Cbl, or SOCS OUTLAST Regulators all showed enhanced tumor cell killing (**Figure 3C**) and IFN-γ and IL-2 cytokine production (**Figure 3D**) compared to CAR T cells without the designs, with most designs showing strong activity even after multiple rounds of tumor co-culture at very low E:T ratios. Under these conditions, the ROR1 CAR alone and the inactivated NR4A TF Modulator negative control completely lost the ability to kill tumor cells and produce cytokine, as expected for T cells that have become dysfunctional due to chronic antigen stimulation (**Figure 3C-D**).

The pan-NR4A OUTLAST Regulators, both the Designed Degrader (SPOP-nr78) and the TF Modulator (Trimer-NR4A-DBD), showed the best tumor cell killing and produced substantially more IFN-γ and IL-2 than the other designs across four independent blood donors (**Figure 3C-D**). The NR4A3-specific Designed Degrader showed reduced killing and cytokine production compared to the pan-NR4A designs across all donors tested, and comparable tumor killing with increased cytokine production relative to the Cbl Designed Degraders tested (**Figure 3C-D**). The Cbl Designed Degraders exemplify the power of our approach to achieve functional benefits with very small cargo sizes; the cbl03-SOCS2box design is less than 400bp of DNA, and this design showed substantial improvements in T cell function over the ROR1 CAR alone (**Figure 3C-D**). The SOCS family Common Substrate Modular design showed enhanced tumor killing comparable to the best pan-NR4A OUTLAST Regulators (**Figure 3C**); however, it showed lower IFN-γ and IL-2 production than the other designs (**Figure 3D, Figure S8C**), as well as higher glucose uptake and more variability in mitochondrial mass (**Figure S9**), indicating that these SOCS family designs may be impacting other aspects of T cell signaling and metabolism in undesired or unexpected ways (EloBC substrate is involved in numerous cell signaling pathways). Based on these results, we decided to advance the best pan-NR4A OUTLAST Regulators for further characterization and testing *in vivo*.

### pan-NR4A designs enhance CAR-T and TCR-T function *in vitro* and *in vivo*

To evaluate the pan-NR4A OUTLAST Regulators in a more therapeutically relevant context, designs were cloned into a single lentiviral vector together with the ROR1 CAR and the EGFRopt safety switch^40^, expressed from either a constitutive MND promoter or an inducible NFkB-derived promoter (**Figure 4A**). We subjected these designs to the serial restimulation with tumor co-culture assay. Across 3 independent blood donors the designs substantially improved tumor killing (**Figure 4B**) and cytokine production (**Figure 4C**) compared to ROR1 CAR alone or ROR1 CAR T cells with a genetic KO of NR4A3. The inducibly-expressed OUTLAST Regulator CAR T cells showed the biggest improvement in performance, with ∼10-fold higher cytokine production than the NRA3 KO CAR T cells after repeated tumor challenge, whereas the constitutively-expressed designs showed ∼3-fold more cytokine production than NR4A3 KO CAR T cells (**Figure 4C**). NR4A activity is known to upregulate the expression of inhibitory receptors PD-1 and TIGIT; flow cytometry following chronic stimulation showed that NR4A OUTLAST Regulators reduced TIGIT and PD-1 expression across both CD4^+^ and CD8^+^ T cell populations (**Figure 4D**). In this experiment, we also included a control that contained only the SPOP degrader domain (no binder); as expected, the ROR1 CAR with SPOP only showed no major changes in function compared to the ROR1 CAR alone (**Figure 4B-D**).

**Figure 4.**
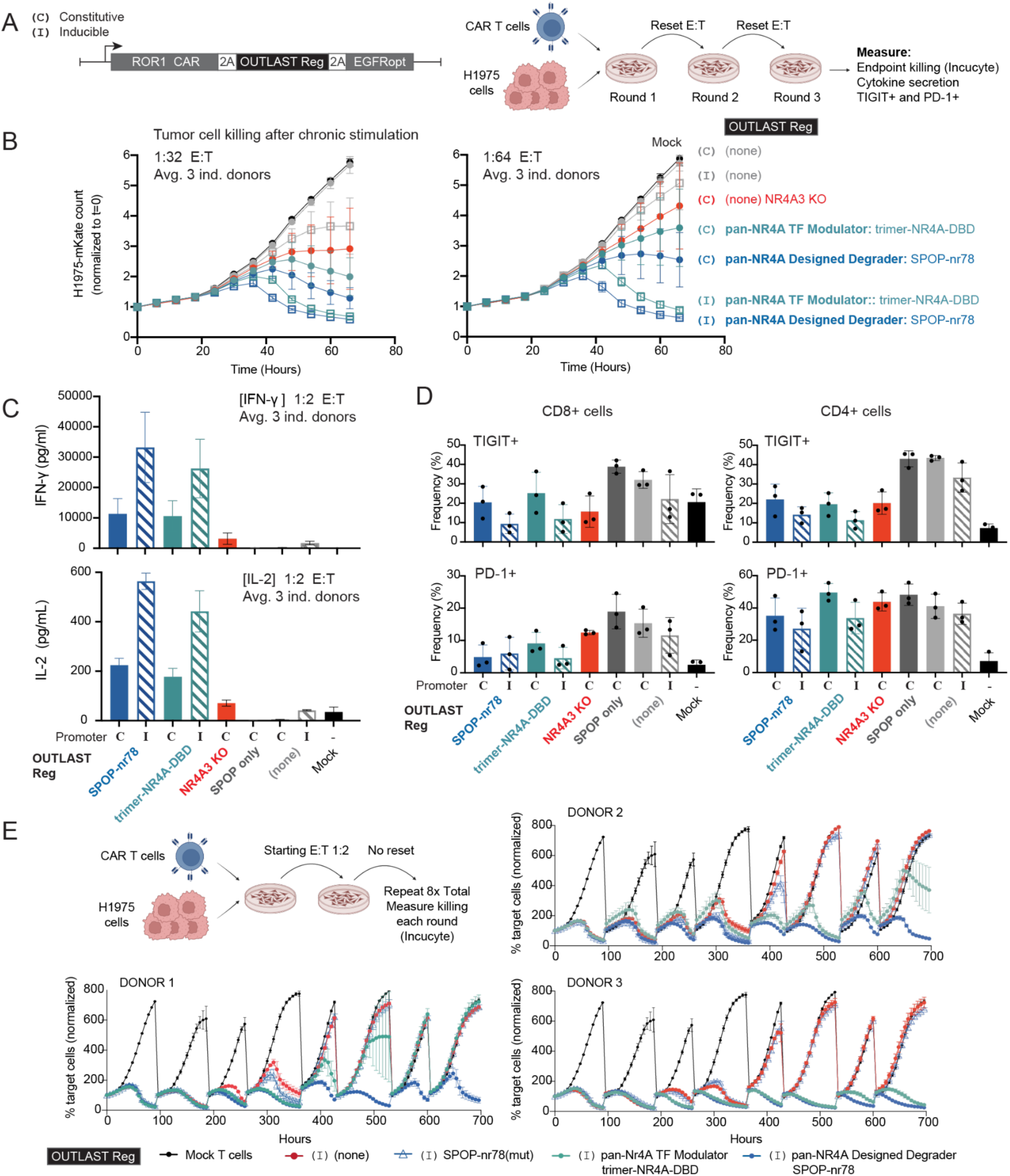
pan-NR4A OUTLAST Regulators improve CAR-T functional persistence and reduce exhaustion markers. (A) OUTLAST Regulator designs were encoded in a single lentiviral vector together with the ROR1 CAR and EGFRopt safety switch under control of a Constitutive MND promoter or an Inducible NFkB-based promoter; CAR T function was assessed by serial restimulation in a tumor co-culture assay. Diagram created using BioRender (https://biorender.com, Fontana, T. 2024). (**B-D**) Data is the average of 3 independent blood donors. NR4A3 KO CAR T cells were generated using CRISPR/Cas9 on a subset of ROR1 CAR-expressing cells. IN/DEL percentages to assess NR4A3 KO efficiency were calculated using ICE analysis (Donor 1: 87 %; Donor 2: 92 %; Donor 3: 77%). (B) Incucyte-based quantification of tumor cell killing kinetics after three rounds of tumor challenge. (C) IFN-γ and IL-2 cytokine secretion measured after three rounds of tumor challenge. (D) TIGIT^+^ and PD-1^+^ frequency assessed by flow cytometry of isolated CD4^+^ and CD8^+^ CAR T cell populations after serial tumor challenge. (E) Incucyte-based quantification of H1975 tumor killing over 8 rounds of serial tumor co-culture challenge; E:T was not reset during the passage of cells in between rounds.

Both the Designed Degrader (SPOP-nr78) and the TF Modulator (timer-NR4A-DBD) showed remarkable enhancement of CAR T function (**Figure 4A-D**). To further differentiate between the two, we performed a more stringent co-culture assay subjecting the CAR T cells to 8 rounds of tumor serial challenge. Even after 8 rounds of co-culture, the Designed Degrader (SPOP-nr78) showed no signs of reduced tumor killing in any of the three donors tested, whereas the TF Modulator started to lose tumor-killing capacity in the final rounds (**Figure 4E**). The binding-ablated SPOP-nr78_mut_ negative control performed similarly to the CAR alone (**Figure 4E**), providing additional support that these designs are functioning via their intended mechanism. To further mechanistically confirm the pan-NR4A Designed Degraders, we also assessed the degradation of endogenous NR4A1 in activated T cells by flow cytometry and intracellular staining with an anti-NR4A1-PE antibody: designs SPOP-nr77, nr77-SPOP, SPOP-nr78, and nr78-SPOP all showed robust degradation (**Figure S10**).

We next evaluated *in vivo* performance of the inducibly-expressed pan-NR4A Designed Degrader (SPOP-nr78) using ROR1^+^ H1975 human xenografts with two independent blood donors and three CAR T cell doses: 2E6, 0.5E6, and 0.1E6 cells (**Figure 5A**). Across both donors and across all three cell doses, ROR1 CAR T cells with the pan-NR4A Designed Degrader (SPOP-nr78) showed 100% survival, a substantial improvement over the inducibly-expressed ROR1 CAR alone and the binding-ablated mutant, SPOP-nr78_mut_ (**Figure 5B**). The ROR1 CAR T cells on their own demonstrated some level of tumor control at the 2E6 dose but not at lower doses, whereas the OUTLAST Regulator CAR T cells were able to control tumors even at the lowest dose of 0.1E6 cells (**Figure 5C**). Furthermore, OUTLAST Regulator CAR T cells exhibited enhanced proliferation (**Figure 5D**) and a more naive/stem-like (CD45RA^+^CD62L^+^) and central memory (CD45RA^-^CD62L^+^) phenotype during the course of the study (**Figure 5E**), properties that are known to correlate with better functional persistence^41^.

**Figure 5.**
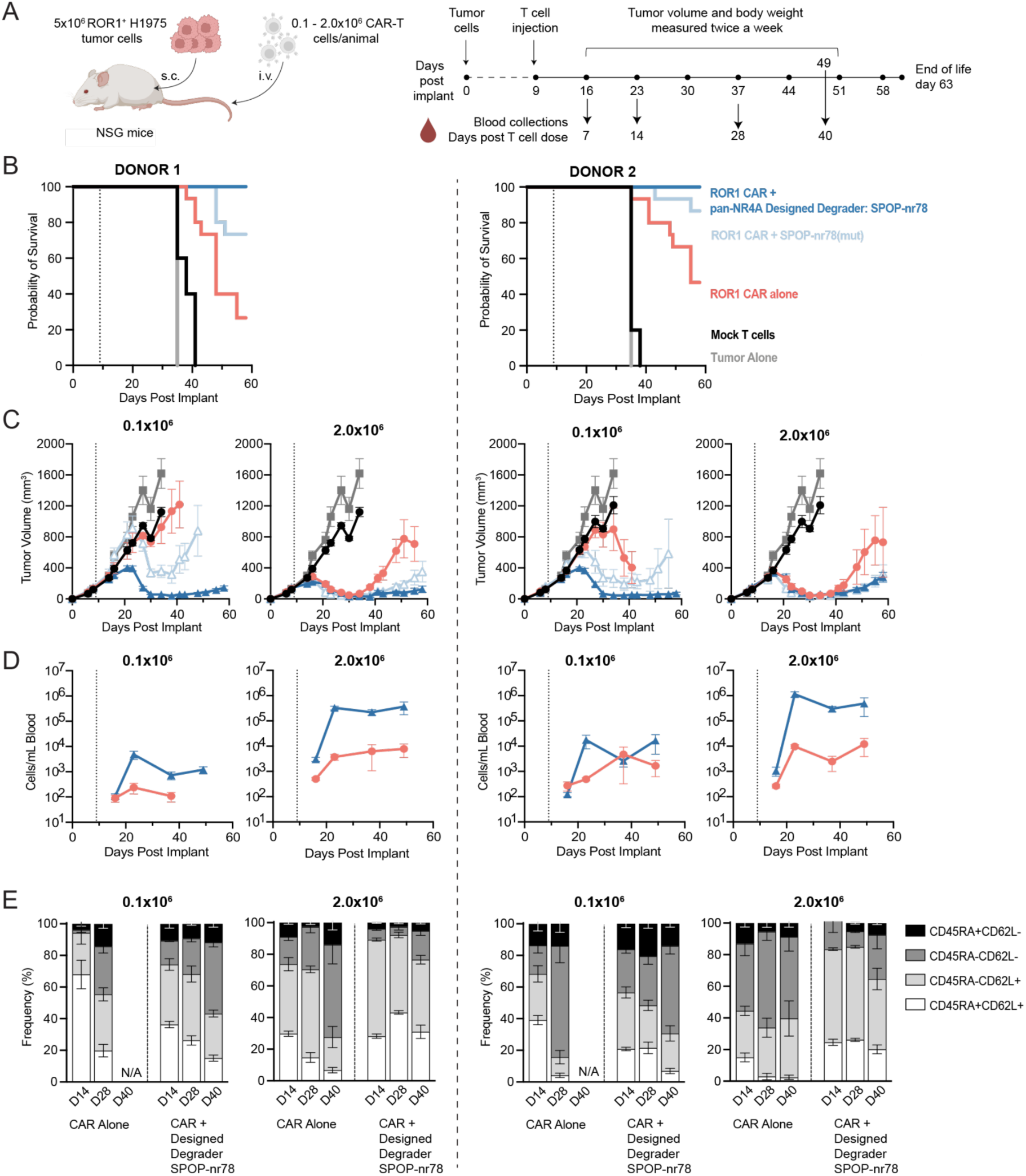
***In vivo* CAR T performance of pan-NR4A Designed Degraders.** (A) Study schematic: ROR1 CAR T cells with or without OUTLAST Regulator designs were administered to H1975 xenografted mice 9 days after tumor implant, with blood collections taken at indicated time points. (B) Survival curves for all mice in the study broken down by human T cell donor. Vertical dotted line indicates CAR T dose in all graphs. For all groups n=5 and average lines stop being plotted with >20% of animals within a group are no longer on study. (C) Average tumor volume over time as measured by calipers at indicated cell doses. (D) Average CAR T cell counts present in peripheral blood over time at indicated cell doses. (E) CAR T cell phenotype measured by flow cytometry assessing CD45RA/CD62L at the indicated days post dose; N/A indicates data not available due to reaching endpoint before samples were collected at that time point.

Lastly, to test the generalizability of the designs beyond CAR T cells, we tested pan-NR4A OUTLAST Regulators for their ability to enhance engineered TCR-T cells. TCRs can target tumor-specific antigens that are challenging for CARs, such as PRAME (PReferentially-Expressed Antigen in MElanoma), a tumor antigen expressed in both solid tumors and hematologic malignancies^42,43^. We tested the function of a clinically-validated PRAME TCR^42^ alone, with the nr77-SPOP pan-NR4A Designed Degrader, or with SPOP alone or nr77 binder alone (negative controls) before or after chronic antigen stimulation with platebound PRAME pMHC, CD58, and ICAM (**Figure 6A**). The Designed Degrader reduced TIGIT expression following chronic stimulation (**Figure 6B**) and improved tumor killing (**Figure 6C**) and IFN-γ and IL-2 cytokine production (**Figure 6D**) when challenged in co-culture with PRAME^+^ H1650 lung adenocarcinoma spheroids. Consistent with results obtained for ROR1 CAR T cells with the designed binders alone (**Figure S11**), PRAME TCR-T cells with the nr77 binder alone also showed intermediate functional improvements (**Figure 6B-D**), suggesting potential inhibition of NR4A function.

**Figure 6.**
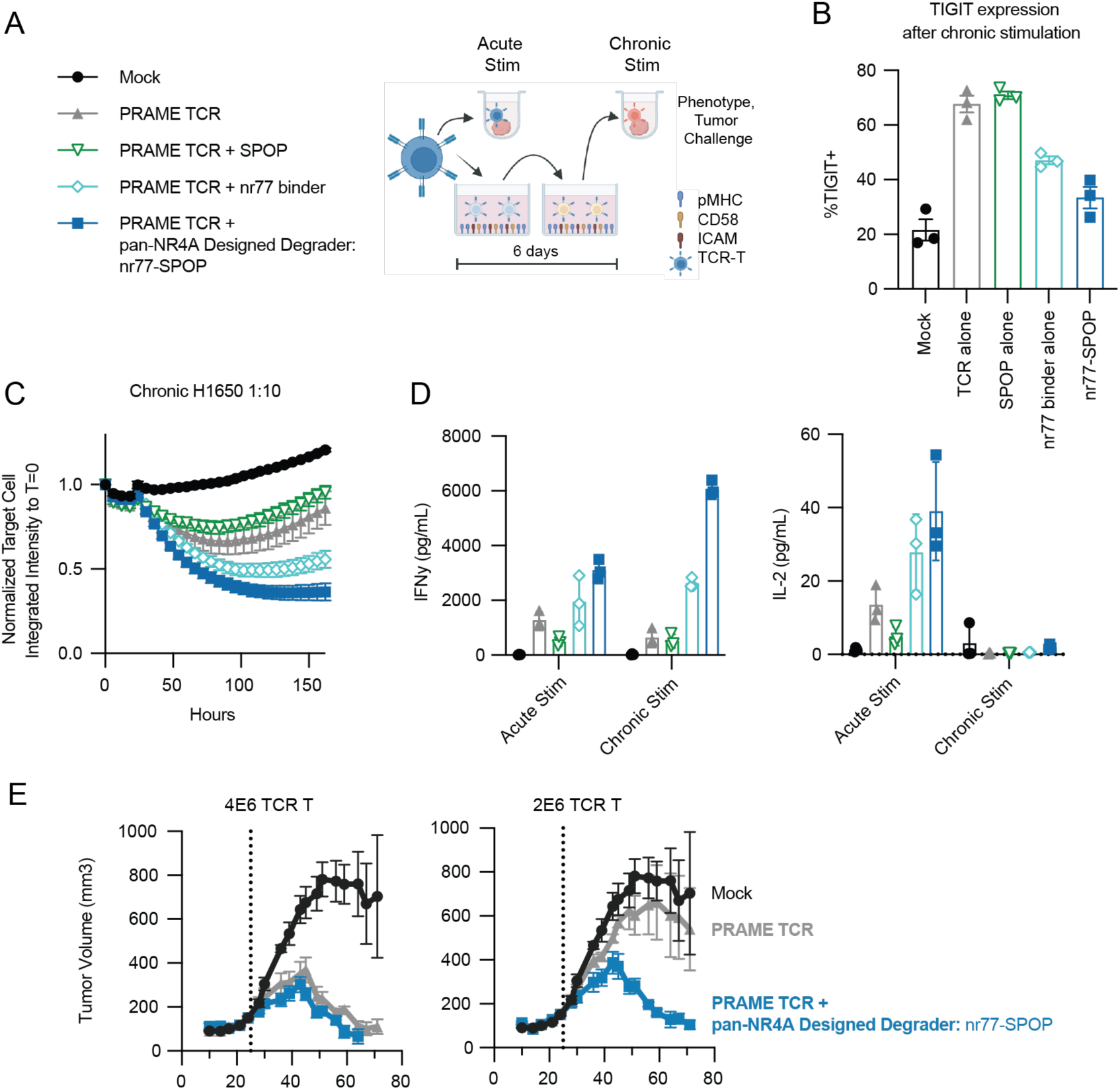
***In vitro* and *in vivo* function of pan-NR4A Designed Degrader in PRAME TCR-T cells.** (A) Constructs tested and schematic of *in vitro* functional assay measuring TCR-T function after chronic stimulation with plate-bound pMHC(PRAME), CD58 and ICAM; experimental diagram created using BioRender (https://biorender.com, Fontana, T. 2024) (B) %TIGIT+ TCR-T cells after chronic stimulation, measured by flow cytometry. (C) Incucyte-based quantification of H1650 tumor killing kinetics after chronic platebound stimulation. (D) IFN-γ and IL-2 cytokine secretion from TCR-T cells measured after chronic platebound stimulation. (E) *In vivo* TCR-T performance measuring average tumor volume in H226-A*02:01 PRAME^+^ xenografted mice at indicated dose levels. Vertical dotted line indicated TCR-T dose; data is the average of 2 independent donors, 5 mice per condition and average lines stop being plotted with >20% of animals within a group are no longer on study.

We then tested the effect of this design on TCR-T efficacy *in vivo* using an additional solid tumor xenograft model with subcutaneously implanted H226-A*02:01 PRAME^+^ lung carcinoma. Engineered PRAME TCR-T cells incorporating the nr77-SPOP OUTLAST Regulator outperformed PRAME TCR-T cells lacking the design, showing robust tumor volume control at cell doses where the TCR-T alone failed (**Figure 6E**). Collectively, these data demonstrate that the pan-NR4A OUTLAST Regulators can dramatically improve engineered T cell function across multiple solid tumor models, both *in vitro* and *in vivo* and in both CAR-T and TCR-T contexts.

## Discussion

Existing cellular engineering efforts have largely taken a one-gene-at-time approach, in which endogenous genes are overexpressed, mutated, deleted or knocked down. Protein design can create new biological functions that transcend the evolutionary constraints of natural proteins. In this study, we focused on well-studied T cell signaling pathways implicated in T cell dysfunction, leveraging computationally-designed protein regulators to simultaneously modulate multiple protein targets. We demonstrated multiple strategies and designs that can enhance the functional persistence of engineered T cells against solid tumors. OUTLAST Regulators can regulate a wide range of targets in a wide range of subcellular locations, from cytoplasmic proteins that act in multiple locations (SOCS family) to ones that primarily act near the cell membrane (Cbl-b and c-Cbl), as well as nuclear-localized transcription factors (NR4A family). The pan-NR4A OUTLAST Regulators performed best overall, dramatically improving both CAR-T and TCR-T function *in vitro* and *in vivo*, with durable tumor control at extremely low cell doses in rigorous solid tumor models (**Figure 5 and 6**).

Achieving pan-family regulation required designing binders against a carefully selected structural epitope, an endeavor uniquely enabled by computational design that would be challenging to achieve with conventional selection-based binder generation techniques. Leveraging AI-based design methods such as RFDiffusion^16^ and ProteinMPNN^17^ coupled with *in silico* analysis to prioritize which designs were tested led to very high success rates: hundreds of our *de novo* NR4A binders achieved low nanomolar binding affinities by yeast display, from which we were able to choose 24 with single-digit nanomolar affinity (**Figure 2A**); all 24 binders exhibited the intended activity in T cells, with a Degradation Score of >0.5 for all three NR4A family members when paired with at least one of the engineered degrader domains (**Figure 2D**), and all of the designs with a Degradation Score >0.5 that were tested in functional assays showed substantial improvements to CAR T cell cytotoxicity and cytokine production (**Figure 2F-G**). These results are notable given that the designs required no wet lab optimization post computational design. Eliminating extensive iteration at the protein design stage enabled us to accelerate the translation of these novel proteins towards creating viable solutions for the complex challenges of cell therapies.

OUTLAST Regulators have the potential to program a broad range of cellular functions across many different therapeutic applications. Their modularity and small cargo footprint allow multiple distinct regulatory components to be stacked within a single engineered cell product. Their heterobifunctional nature can couple the simultaneous binding of multiple targets with diverse biological outputs, ranging from proteasomal degradation to control of gene expression to modulation of signal transduction. While this work used *ex vivo*-edited T cells, OUTLAST Regulators could readily be incorporated into other cell types and gene therapy modalities. Unlike static gene knockouts or constitutive overexpression, the OUTLAST Regulator platform opens the door to dynamic regulation of cellular functions, where multiple designs can be combined under the control of regulated promoters and context-dependent stimuli, facilitating the engineering of complex biological circuitry.

## Acknowledgments

We thank Stan Riddell and Margo Roberts for helpful discussions and feedback on the manuscript. We thank Lindsie Goss for helpful feedback on the manuscript, and Andrew Ng for helpful discussions during the early stages of this project; Kristie Shirley and Lesley Jones for cloning support; Allan Wang and Venessa Montoya for lentivirus production support; Leah Tait, Jared Hammer, and Maria Steele for *in vitro* assay support and assistance with data analysis; John Crowl and Chelsea Dunmire for assistance with gene KO studies; Robert Lenox-Pulgarin and Aye Chen for assistance with the monobody library; Jerry C. Chen for technical support with *in vivo* studies, and David Clausen for statistical analysis support.

## Author Contributions

S.E.B. and M.J.L. conceived of the study. B.C. performed computational protein design for *de novo* binders against NR4A and Cbl with input from S.E.B., B.D.W., and R.A.L., who selected which designs were advanced for functional testing. S.M., F.H., H.F.M., and S.E.B. designed the degradation domains and Designed Degrader constructs; D.N. contributed to engineering and screening degrader domains. G.W.F. contributed to initial design work for the NR4A binders and Designed Degraders. S.E.B., R.A.L., and T.M.D. designed the TF Modulator proteins, S.E.B. designed the Common Substrate Modulator proteins, and F.H. and S.M. designed the constructs. J.D. developed the NR4A3-specific monobody binder. I.G. performed yeast surface display experiments. B.H. performed protein purification and biophysical characterization and B.H., T.M.D., and S.Y. analyzed binding kinetics data. F.H., S.M., and J.C. designed and performed target degradation assays and analyzed the data. S.M., J.C., and F.H. performed *in vitro* CAR T cell functional assays and analyzed data; H.F.M. and R.A. designed experiments and analyzed data. K.G.H. and designed and performed *in vivo* studies and analyzed the data. S.E.B., S.M., R.A.L., H.F.M., and K.G.H. visualized data and generated the figures. S.E.B. and M.J.L. wrote the paper; all authors reviewed and edited the manuscript. M.J.L, D.B., A.E.F., and S.E.B. supervised the work.

## Declaration of Interests

Outpace Bio and the University of Washington have filed patent applications related to this work.

## Declaration of AI-assisted technologies in the writing process

The authors used Claude (Anthropic) and Gemini (Google) to assist with grammar and sentence structure in this manuscript. After using these tools, authors reviewed and edited the content as needed. The authors take full responsibility for the content of the publication.

---

**Figure S1. (related to Fig. 1).**
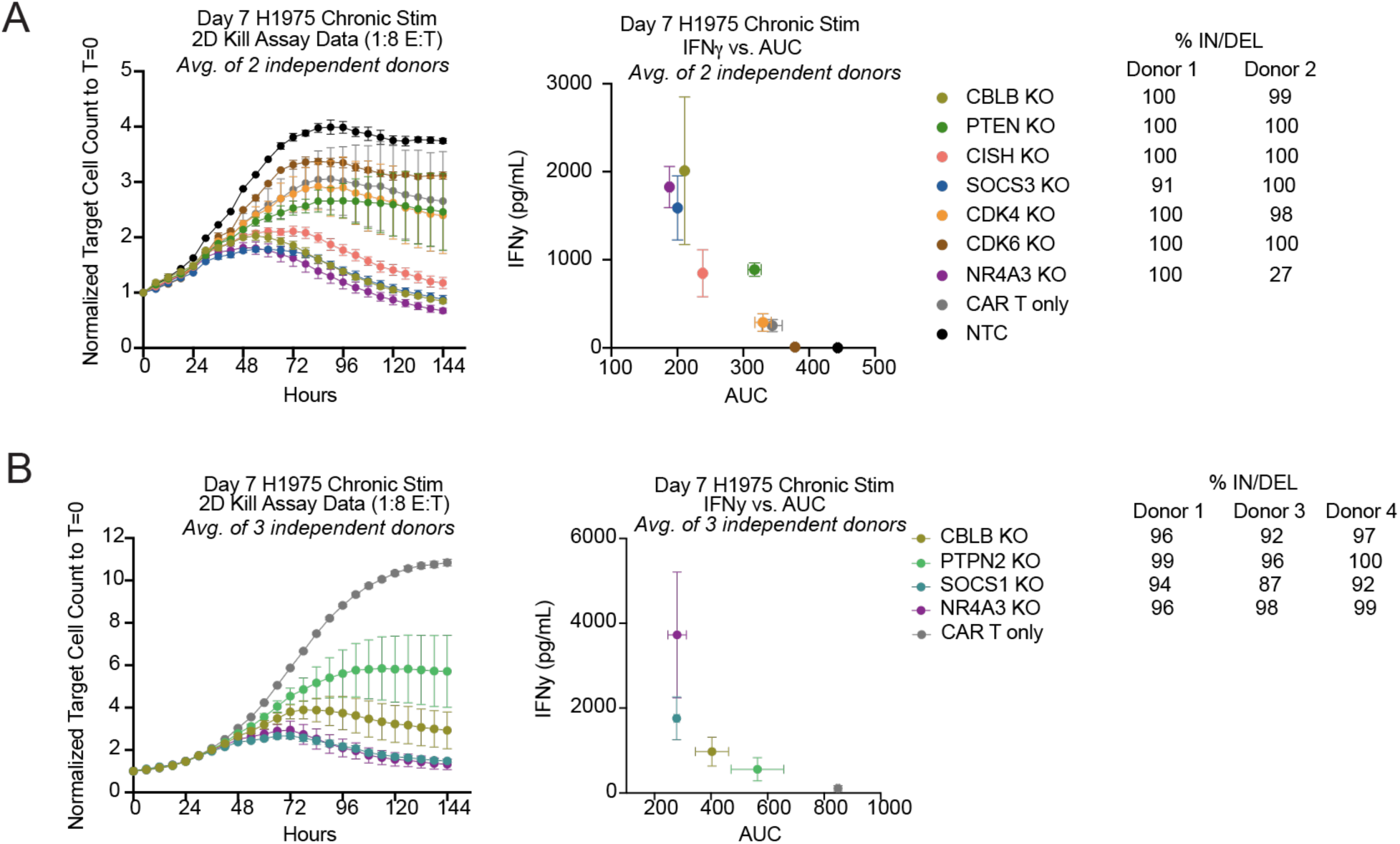
***In vitro* functional assessment of ROR1 CAR T cells with genetic knockouts (KOs) after chronic stimulation against ROR1^+^ H1975 tumor cell line.** (**A-B**) Incucyte-based quantification of H1975 tumor cell killing after three rounds of tumor co-culture challenge (*left*), and a plot of area under the curve (AUC) of tumor killing plot versus IFN-γ (*right*); lower AUC and higher IFN-γ indicates better CAR T cell performance. Data shown is the average of two independent blood donors; the percentage insertion/deletion (%IN/DEL) for the gene KOs for each donor is indicated. The data in (A) and (B) are from two separate experiments in which different sets of gene KOs were tested.

**Figure S2. (related to Fig. 1).**
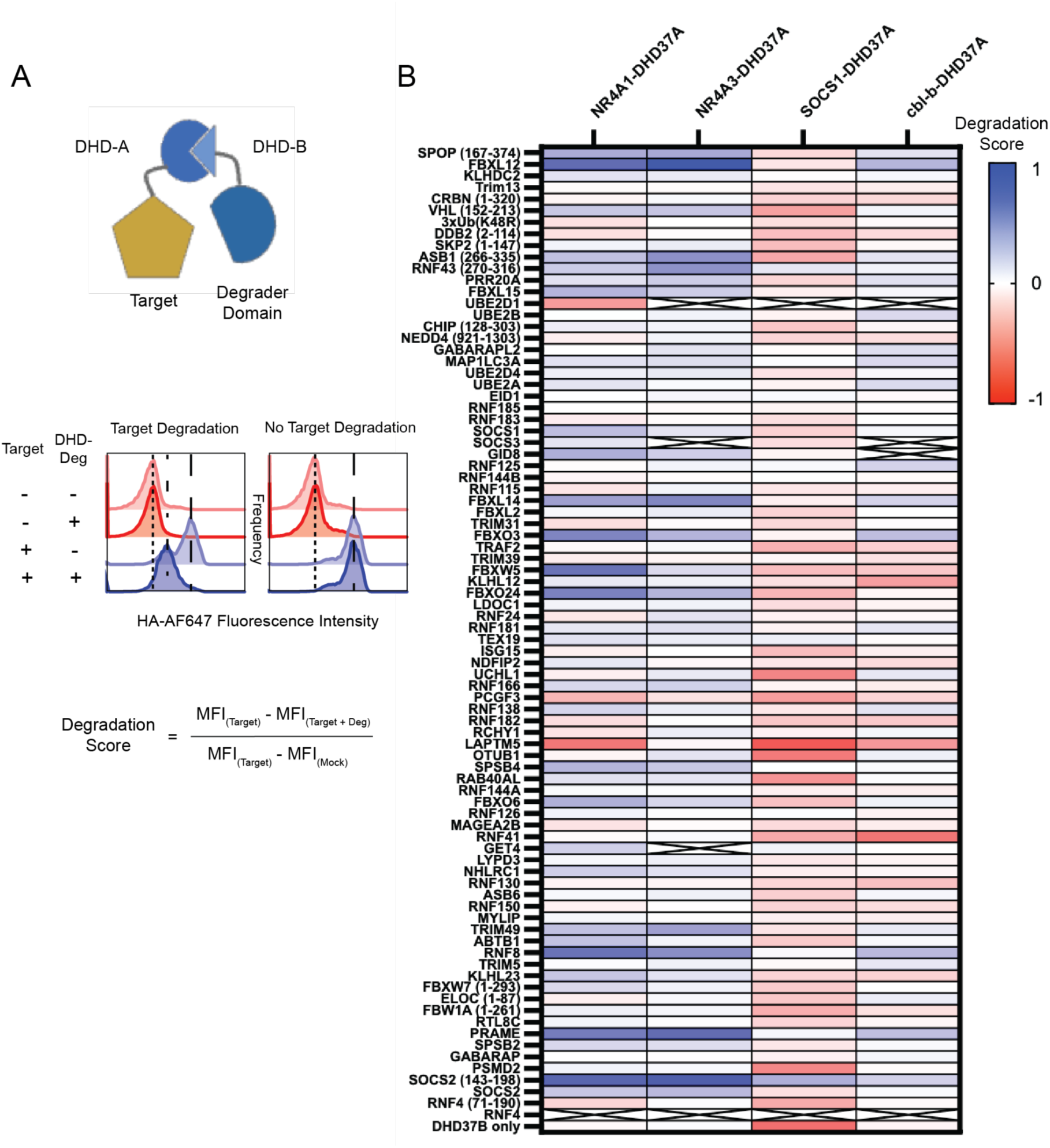
Screen of degrader domains capable of down-regulating targets of interest in primary human T cells. **(A)** Schematic of the designed heterodimer (DHD)–degrader screening strategy. HA-tagged protein targets of interest, NR4A1, NR4A3, SOCS1, and Cbl-b, were each fused to one component of a DHD, while a library of candidate E3 ligase degrader domains was fused to the complementary DHD component. Pairwise co-expression of these components enabled systematic assessment of target degradation, with HA-tag levels quantified as a readout in the presence or absence of each degrader construct. “Degradation Score” was derived from mean fluorescence intensity (MFI) values using the formula shown; a Degradation Score of 1.0 indicates complete degradation. **(B)** Degradation Score heatmap for all degrader domains evaluated against each target of interest. An “X” indicates that data was not collected for that pair.

**Figure S3. (related to Fig 2).**
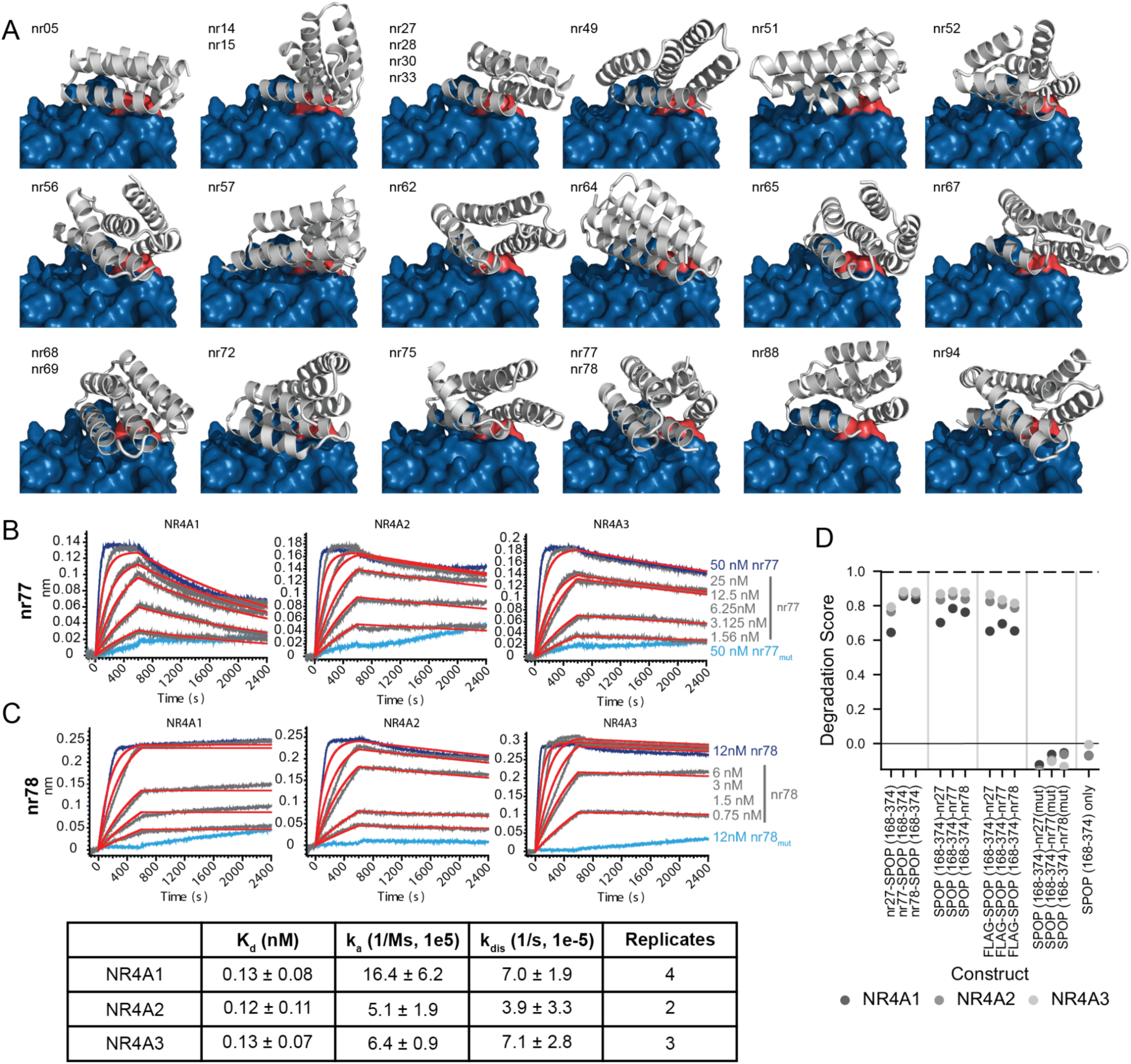
Characterization of *de novo* pan-NR4A binders. (**A)** AlphaFold2-predicted structural models of designed binders (gray cartoon representation) in complex with NR4A1 (PDB: 3V3E, blue surface). The conserved C-terminal LBD motif is highlighted in coral. Designs with unique sequences on the same backbone are indicated by multiple labels per binder image. (**B–C**) Biolayer interferometry (BLI) sensorgrams measuring binding of biotinylated NR4A1, NR4A2, and NR4A3 target proteins captured on streptavidin biosensors to nr77 over a 2-fold dilution series from 50 nM to 1.56 nM (B) and nr78 over a two-fold dilution series from 12 nM to 0.75 nM (C). Binders with rational mutations to impair key binding interface residues, nr77_mut_ and nr78_mut_, showed no detectable binding at 50 nM for nr77 or 12 nM for nr78 (cyan). Estimated dissociation constants (K_d_) for nr78 were derived from 1:1 global kinetic fits of the sensorgram data in (C). (**D**) Degradation Score (see Methods, Figure 2) of overexpressed NR4A1/2/3 target protein in primary human T cells in response to Designed Degrader designs assessing effect of SPOP fusion orientations (binder–SPOP vs SPOP–binder) and tag (FLAG–ligase–binder); SPOP only control and binding interface mutations show no degradation activity.

**Figure S4. (related to Fig 2).**
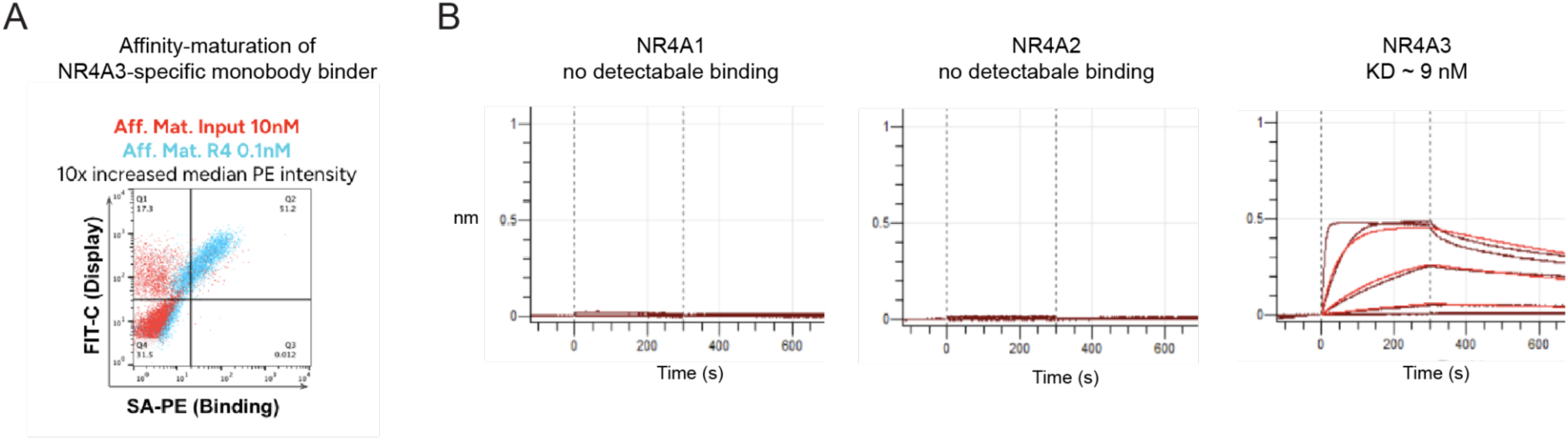
Development of an NR4A3-specific monobody. (A) Representative FACS plots from the anti-NR4A3 monobody affinity maturation. The BC, DE, and FG loops were amplified by error-prone PCR (GeneMorph II, Agilent) from polyclonal phagemid DNA recovered after four rounds of panning a 2.95 × 10¹⁰-member synthetic FN3 library (UniProt P02751, aa 1538–1631, D1540S) against NR4A3-LBD, shuffled, and cloned into pETCONV3 for display in EBY100 (red). The library was sorted against NR4A3-LBD for four rounds (blue). Sanger sequencing of the output identified clones for kinetic and biophysical characterization. (B) Biolayer interferometry (BLI) sensorgrams measuring binding of biotinylated NR4A1, NR4A2, and NR4A3 target proteins captured on streptavidin biosensors to the monobody analyte in a 4-fold dilution series from 1000 nM to 0.98 nM; dissociation constant (K_d_) for NR4A3 was derived from 1:1 global kinetic fitting of BLI sensorgram data.

**Figure S5. (related to Fig. 2).**
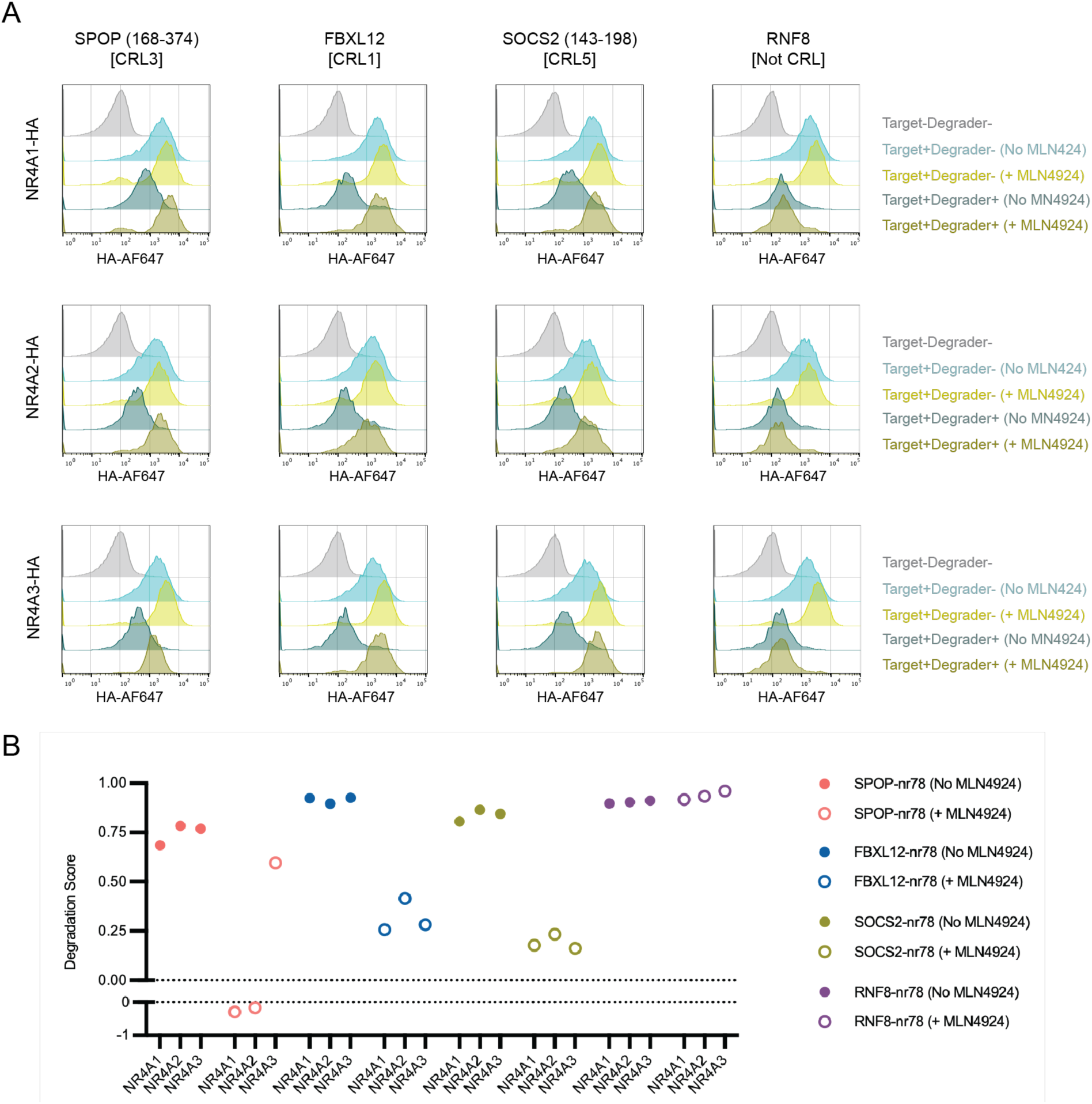
Designed Degraders with Cullin-RING Ligase (CRL)-mediated degrader domains function via expected proteasomal degradation pathways. (A) Flow cytometry data monitoring degradation of HA-tagged NR4A1 in the presence of absence of MLN4924 for indicated Degrader Domains fused to nr78 binder. (B) Degradation Score derived from flow cytometry data (A).

**Figure S6. (related to Fig. 1 and 3).**
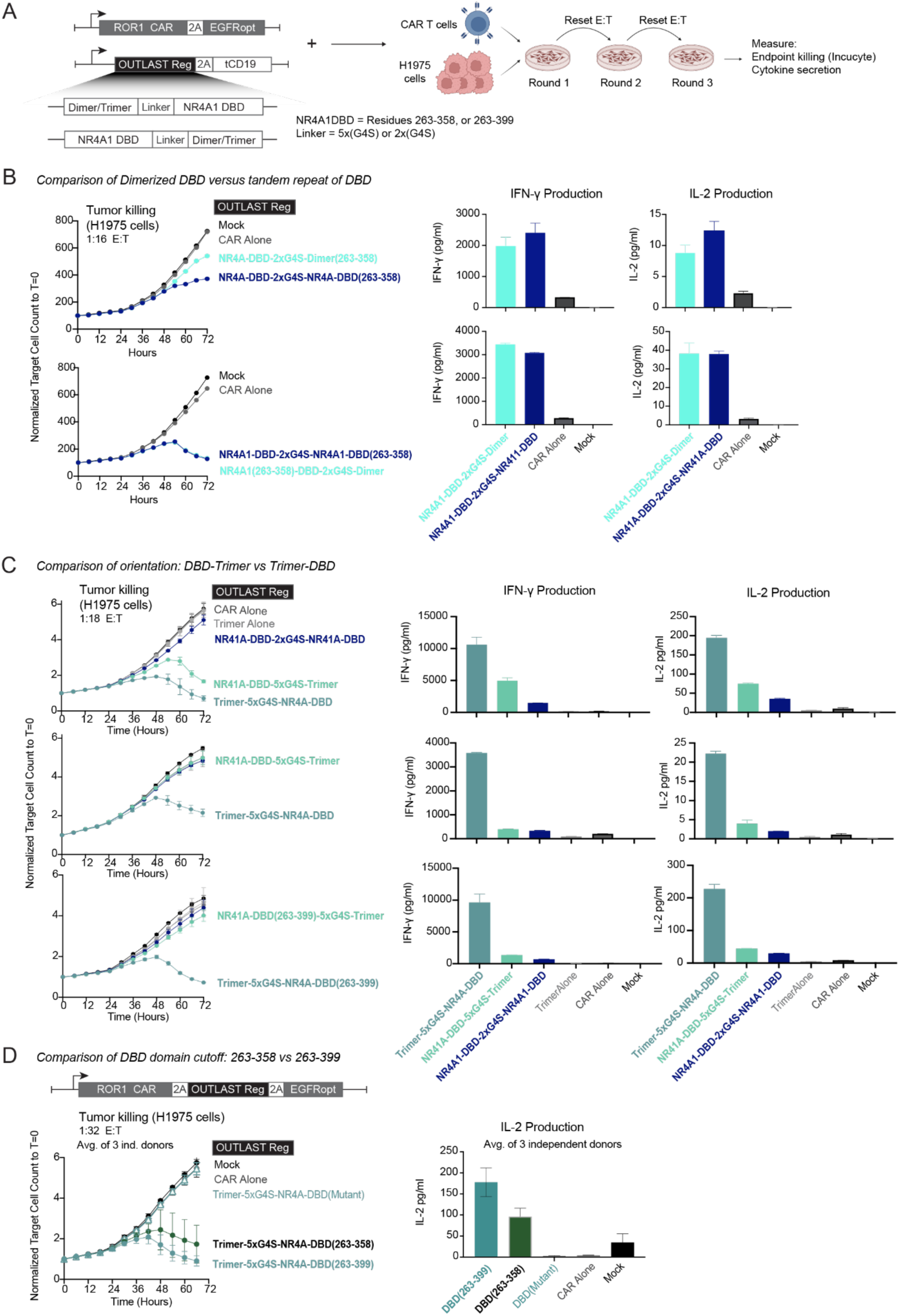
Characterization of pan-NR4A TF Modulator designs. (A) Design of pan-NR4A TF Modulator constructs: The DNA-binding domain (DBD) of NR4A1 was fused to *de novo* designed homodimerizing and homotrimerizing domains. (**B-D**) Incucyte-based quantification of H1975 tumor cell killing kinetics after three rounds tumor co-culture challenge (*left)*; IFN-γ and IL-2 cytokine secretion measured at Day 7 after three rounds of tumor challenge (*right*). (B) Comparison of Dimer design to a tandem repeat of the DBD as a control; data is shown for two independent blood donors. (C) Comparison of orientation: fusing the trimerization domain to N-terminus vs C-terminus of the DBD; data is shown for three independent blood donors. (D) Comparison of a shorter (NR4A1 residues 263-358) and longer (NR4A1 residues 263-399) DBD; data is the average of 3 independent blood donors.

**Figure S7. (related to Fig. 1 and 3).**
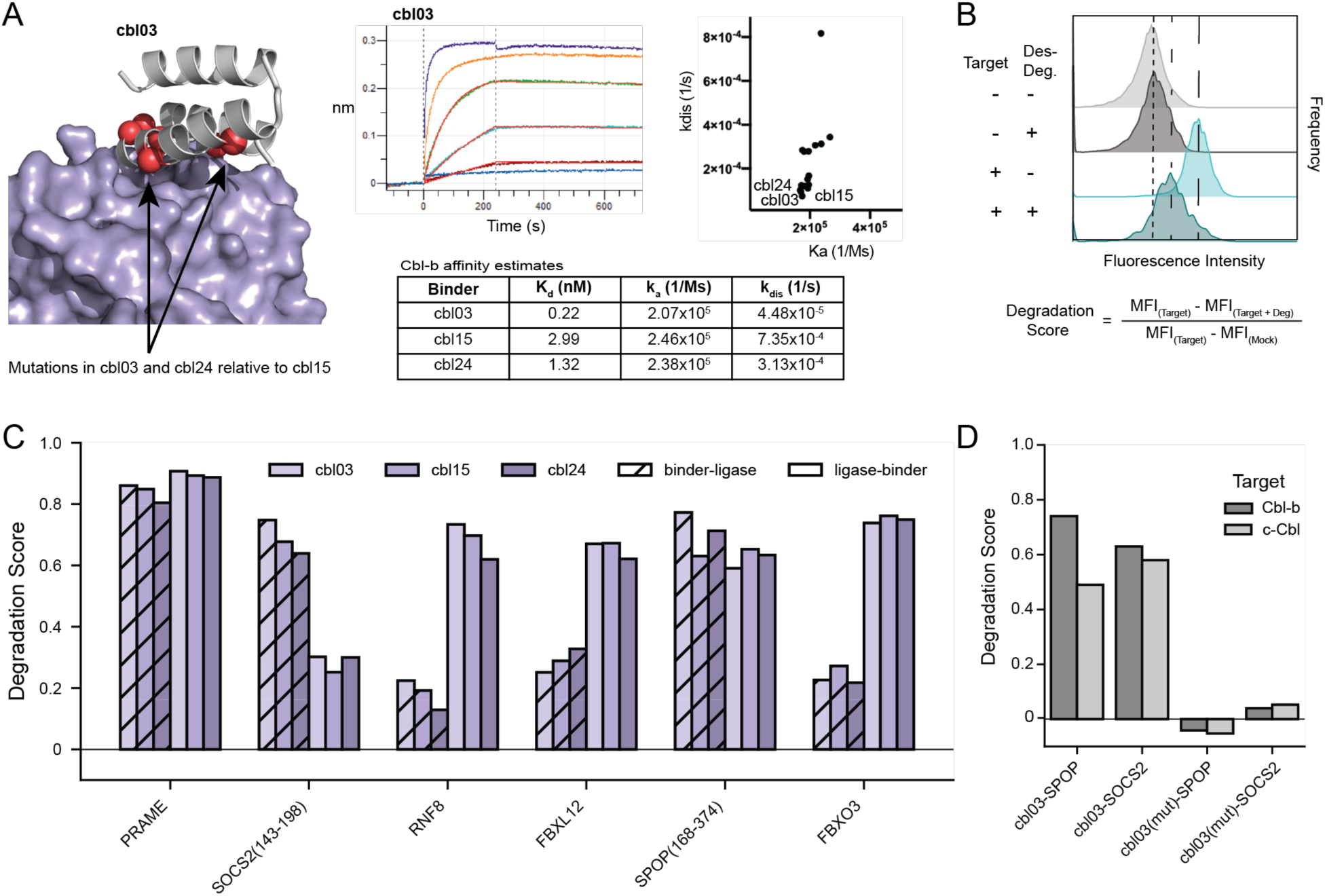
Characterization of Cbl Designed Degrader designs. (A) AlphaFold2-predicted structural models of representative designed binders (gray cartoon representation) in complex with Cbl-b (PDB: 3VGO, purple surface); all three of the best binders share a common backbone architecture; positions that differ among the three variants are indicated by coral spheres. Designs were assessed for binding to biotinylated Clb-b target proteins by BLI, and dissociation constant (K_d_) estimates were derived from global 1:1 kinetic fits of the sensorgrams. (B) Representative histogram from the target degradation assay, in which the target protein (Cbl-b or c-Cbl) is overexpressed in the presence or absence of the binder-degrader fusion. Degradation Score is calculated according to the equation shown. (C) Degradation Scores for Cbl-b Target protein for the three binders (depicted in shades of purple) fused to six distinct Degrader Domains (x-axis) in two different fusion orientations: binder–ligase (solid) and ligase–binder (hatched). (D) Degradation Scores for Cbl-b and c-Cbl Target protein in primary human T cells for Designed Degrader designs fusing cbl03 binder to either SPOP(168-374) or SOCS2(143-198) Degrader Domains, compared against binding-deficient variants harboring interface mutations that ablate Target binding.

**Figure S8. (related to Fig. 1 and 3).**
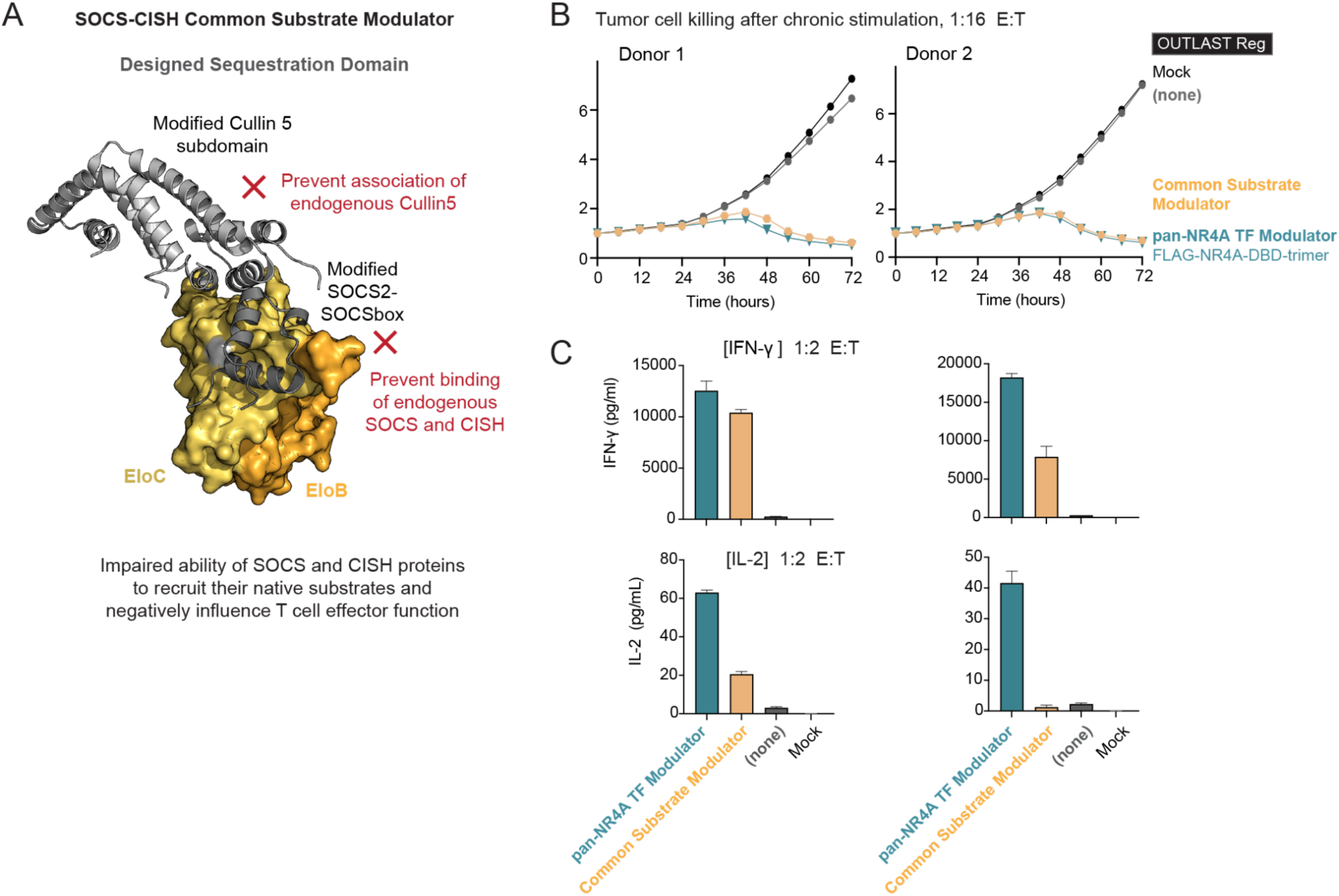
***In vitro* CAR T performance of the SOCS family Common Substrate Modulator designs.** (A) Structural model and intended mechanism for the SOCS-CISH Common Substrate Modulator design. (B) Incucyte-based quantification of H1975 tumor cell killing kinetics after three rounds of tumor challenge. (C) IFN-γ and IL-2 cytokine secretion measured after three rounds of tumor challenge; the TF Modulator design “FLAG-NR4A-DBD-trimer” was included as a positive control.

**Figure S9. (related to Fig. 3).**
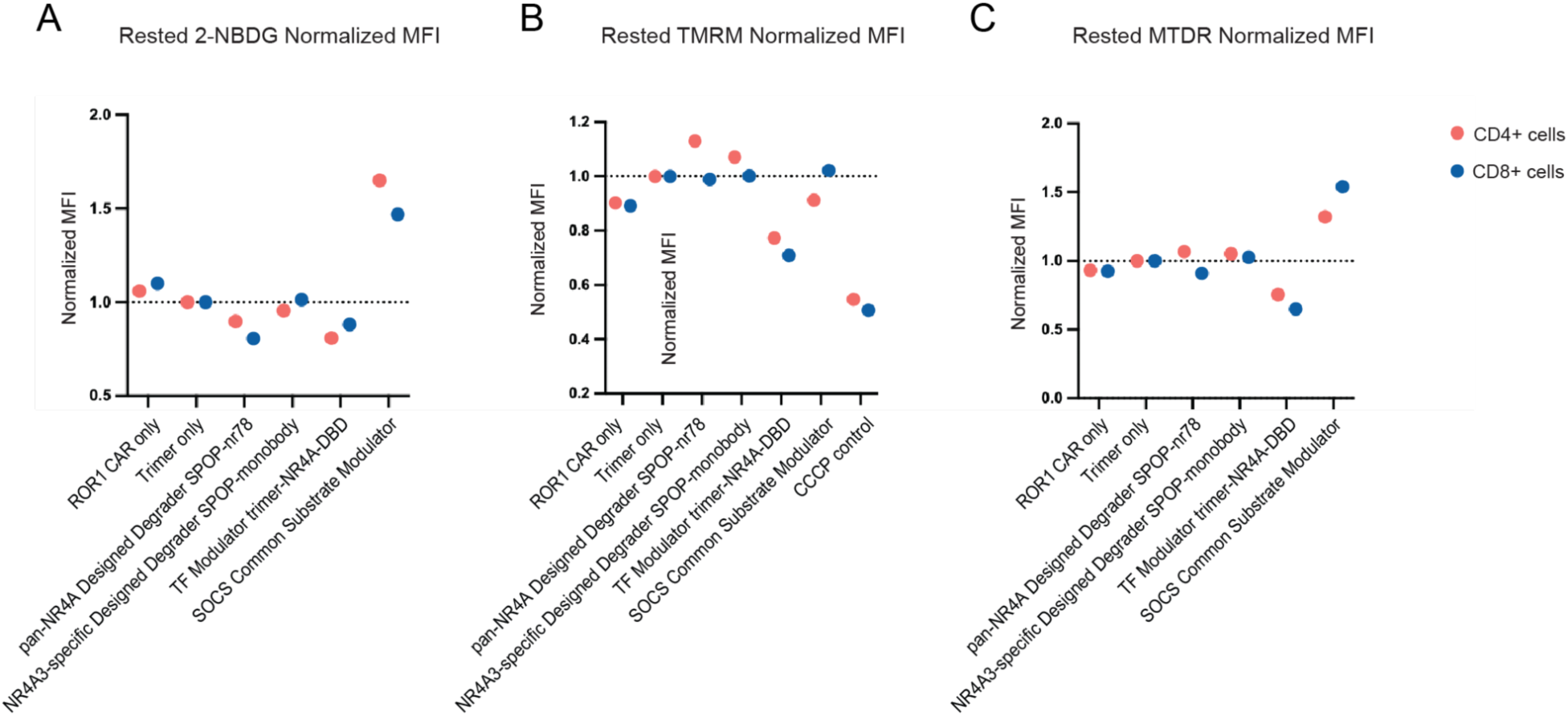
Glucose uptake and mitochondrial membrane potential and mass for OUTLAST Regulator CAR T cells. (A) Glucose uptake, measured by flow cytometry MFI of NBDG (a fluorescent glucose analog) to monitor cellular glucose uptake. (B) Mitochondrial membrane potential, measured by flow cytometry MFI of tetramethylrhodamine methyl ester (TMRM). (C) Mitochondrial mass, measured by flow cytometry MFI of MitoTracker Deep Red (MTDR).

**Figure S10. (related to Fig. 4).**
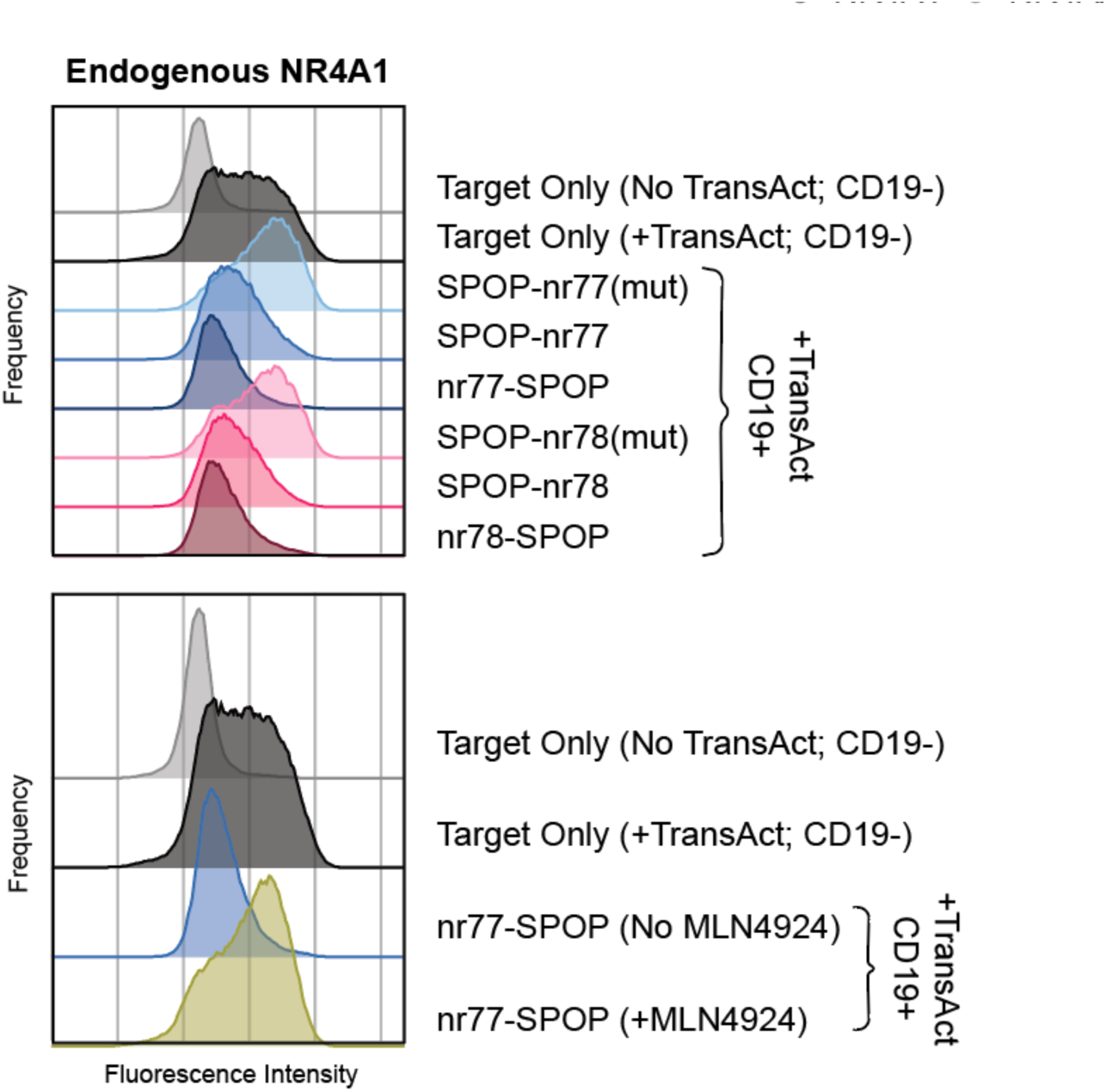
Degradation of endogenous NR4A1 Target in primary T cells. Degradation of endogenous NR4A1 target protein in primary human T cells, in response to the transduced pan-NR4A Designed Degrader designs indicated; intracellular staining for endogenous NR4A1 was performed by incubating cells with anti-NR4A1-PE antibody (see **Methods**); T cells transduced with Designed Degrader design constructs were treated with TransAct beads to activate the T cells and stimulate NR4A expression.

## METHODS

### RESOURCE AVAILABILITY

#### Lead contact

Requests for further information and resources should be directed to the lead contact, Scott E. Boyken.

#### Materials Availability

Plasmids generated in this study can be made available on request for non-commercial use. Materials from this study for potential commercial use can potentially be made available with a completed Material Transfer Agreement.

#### Data and Code Availability

● The Rosetta macromolecular modelling suite (https://www.rosettacommons.org) is freely available to academic and non-commercial users. Commercial licences for the suite are available through the University of Washington Technology Transfer Office.
● RFDiffusion code can be downloaded from https://github.com/RosettaCommons/RFdiffusion
● ProteinMPNN code is available from https://github.com/dauparas/ProteinMPNN and https://github.com/sokrypton/ColabDesign/
● AlphaFold2 is available from https://github.com/google-deepmind/alphafold
● OpenFold code is available from https://github.com/aqlaboratory/openfold
● Any additional information required to reanalyze the data reported in this paper is available from the lead contact upon request.

### Key Resources Table

| REAGENT OR RESOURCE | SOURCE | IDENTIFIER |
| --- | --- | --- |
| Antibodies |  |  |
| FITC anti-human CD19 | BioLegend | Cat# 302206;<br>RRID:AB_314236 |
| BV421 anti-human EGFR | BioLegend | Cat# 352911;<br>RRID:AB_2562213 |
| Alexa Fluor 647 anti-HA-Tag | Cell Signaling Technology | Cat# 3444s;<br>RRID:AB_10693329 |
| PE anti-DYKDDDDK (FLAG) Tag | BioLegend | Cat# 637310;<br>RRID:AB_2563148 |
| BUV395 anti-human CD8a | BD Biosciences | Cat# 569178;<br>RRID:AB_3684844 |
| BUV496 anti-human CD4 | BD Biosciences | Cat# 612936;<br>RRID:AB_2870220 |
| PE anti-human NR4A1 | Thermo Fisher Scientific | Cat# 12-5965-82;<br>RRID:AB_1257209 |
| BV421 anti-DYKDDDDK (FLAG) Tag | BioLegend | Cat# 637322;<br>RRID:AB_2750061 |
| APC anti-human CD19 | BioLegend | Cat# 302212;<br>RRID:AB_314242 |
| Recombinant human ROR1-Fc conjugated to Alexa Fluor 647 | Outpace Bio | Custom reagent |
| BV421 anti-human CD8a | BD Biosciences | Cat# 740078;<br>RRID:AB_2739841 |
| Alexa Fluor 488 anti-human EGFR | BioLegend | Cat# 352908;<br>RRID:AB_11126165 |
| PE anti-human TIGIT | BioLegend | Cat# 372704;<br>RRID:AB_2632730 |
| PE-Cy7 anti-human PD-1 | Thermo Fisher Scientific | Cat# 25-2799-42;<br>RRID:AB_10853804 |
| BUV395 anti-human CD45RA | BD Biosciences | Cat# 568712;<br>RRID:AB_3684488 |
| BUV615 anti-human CD62L | Thermo Fisher Scientific | Cat# 366-0629-41;<br>RRID:AB_2925411 |
| BUV737 anti-human CD27 | BD Biosciences | Cat# 612829;<br>RRID:AB_2870151 |
| BV711 anti-human CD28 | BioLegend | Cat# 302948;<br>RRID:AB_2616857 |
| Super Bright 780 anti-human LAG3 | Thermo Fisher Scientific | Cat# 78-2239-42;<br>RRID:AB_2802471 |
| APC anti-human TIM3 | BioLegend | Cat# 364804;<br>RRID:AB_2910409 |
| BV510 anti-human CD39 | BioLegend | Cat# 328219;<br>RRID:AB_2563265 |
| BV650 anti-human CD3 | BioLegend | Cat# 317324;<br>RRID:AB_2563352 |
| BV785 anti-human CD8 | BioLegend | Cat# 301046;<br>RRID:AB_2563264 |
| APC anti-human hCD45 | BioLegend | Cat# 368512;<br>RRID:AB_2566372 |
| BV421 anti-human CD19 | BioLegend | Cat# 302234;<br>RRID:AB_11142678 |
| BV711 anti-human EGFR | BioLegend | Cat# 352920;<br>RRID:AB_2687123 |
| BUV395 anti-human CD8 | BD Biosciences | Cat# 563795;<br>RRID:AB_2722501 |
| BUV805 anti-human CD3 | BD Biosciences | Cat# 612893;<br>RRID:AB_2870181 |
| PE anti-human CD56 (NCAM) | BioLegend | Cat# 304606;<br>RRID:AB_314448 |
| Biotin anti-human EGFR Antibody | BioLegend | Cat# 352934;<br>RRID:AB_2922568 |
| BV421 anti-human HLA-A2 | BioLegend | Cat# 343326;<br>RRID:AB_2721523 |
| APC anti-human TCR Vb8 | BioLegend | Cat# 348106;<br>RRID:AB_2721307 |
| SULFO-TAG anti-hu IFNg antibody | Meso Scale<br>Discovery | Cat# D21Q0-3 |
| SULFO-TAG anti-hu IL-2 antibody | Meso Scale<br>Discovery | Cat# D21QQ-3 |
| Critical Commercial Assays |  |  |
| V-plex Proinflammatory Panel 1 Human Kit | Meso Scale<br>Discovery | Cat# K15049D-4 |
| Quick-DNA MiniPrep w/Zymo-Spin IIC Columns,<br>Capped, 200 Preps | Research<br>Products<br>International Corp | Cat# 50-197-7251 |
| DNA Clean & Concentrator-5 w/ Zymo-Spin IC<br>Columns | Research<br>Products<br>International Corp | Cat# 50-197-7310 |
| P3 Primary Cell 4D-Nucleofector X Kit S | Lonza | Cat# NC0545312 |
| EasySep Human T Cell Isolation Kit | Stemcell<br>Technologies | Cat# 100-0695 |
| EasySep Human CD19 Positive Selection Kit II | Stemcell<br>Technologies | Cat# 17854 |
| Chemicals, peptides, and recombinant proteins |  |  |
| MLN4924 (Pevonedistat) | Selleckchem | Cat# S7109 |
| Recombinant Human IL-2 Protein | R&D Systems | Cat# 202-IL |
| Recombinant Human IL-7 Protein | R&D Systems | Cat# 207-IL |
| Recombinant Human IL-15 Protein | R&D Systems | Cat# 247-ILB |
| Biotinylated PRAME pMHC | Outpace Bio | Custom reagent |
| Recombinant Human CD58/LFA-3 Fc Chimera Protein, CF | R&D Systems | Cat# 10068-CD |
| Recombinant Human ICAM-1/CD54 Fc Chimera Protein, CF | R&D Systems | Cat# 720-IC |
| NeutrAvidin Protein | ThermoFisher | Cat# 31000 |
| CTS OpTmizer T Cell Expansion SFM, bottle format | Thermo Fisher Scientific | Cat# A1048501 |
| Immune Cell Serum Replacement (CTS SR) | Thermo Fisher Scientific | Cat# A2596102 |
| L-Glutamine 200mM | Thermo Fisher Scientific | Cat# 25030081 |
| CTS GlutaMAX Supplement | Thermo Fisher Scientific | Cat# A1286001 |
| CryoStor CS10 | Stemcell Technologies | Cat# 7930 |
| RPMI 1640 Medium, GlutaMAX Supplement | Thermo Fisher Scientific | Cat# 61870036 |
| Fetal Bovine Serum, certified, heat inactivated, United States | Thermo Fisher Scientific | Cat# 10082147 |
| TrypLE Express Enzyme (1X), phenol red | Thermo Fisher Scientific | Cat# 12605036 |
| Hygromycin B (50 mg/mL) | Thermo Fisher Scientific | Cat# 10687010 |
| Blasticidin (10 mg/mL) | Thermo Fisher Scientific | Cat# A11139-03 |
| Puromycin Dihydrochloride | Thermo Fisher Scientific | Cat# A1113803 |
| T Cell TransAct, human | Miltenyi | Cat# 130-111-160 |
| ViaStain AOPI Staining Solution, 5 mL | Revvity | Cat# CS2-0106-5ML |
| PBS, pH 7.2 | Thermo Fisher Scientific | Cat# 20012043 |
| Matrigel | Corning | Cat# 356231 |
| 20X PBS Tween | Thermo Fisher Scientific | Cat# 28352 |
| TrueCut Cas9 Protein v2 | Thermo Fisher Scientific | Cat# A36499 |
| Q5 High-Fidelity 2X Master Mix | New England Biolabs | Cat# NC0355749 |
| MojoSort Streptavidin Nanobeads | BioLegend | Cat# 480016 |
| MojoSort Buffer (5X) | BioLegend | Cat# 480017 |
| EasySep Buffer | Stemcell Technologies | Cat# 20144 |
| Brilliant Stain Buffer | BD Biosciences | Cat# 563794 |
| Cell Staining Buffer | BioLegend | Cat# 420201 |
| Fluorofix Buffer | BioLegend | Cat# 422101 |
| Foxp3 / Transcription Factor Staining Buffer Set | Thermo Fisher Scientific | Cat# 00-5523-00 |
| UltraComp eBeads Plus Compensation Beads | Thermo Fisher Scientific | Cat# 01-3333-41 |
| Precision Count Beads | BioLegend | Cat# 424902 |
| RBC Lysis/Fixation Solution (10X) | BioLegend | Cat# 422401 |
| HLA-A*0201 (SLLQHLIGL), PE (PRAME pMHC Dextramer-PE) | Immudex | Cat# WB04074-PE |
| eBioscience™ Fixable Viability Dye eFluor™ 780 (FVD780) | Thermo Fisher Scientific | Cat# 65-0865-18 |
| 2-NBDG | Thermo Fisher Scientific | Cat# N13195 |
| MitoTracker™ Deep Red FM Dye, for flow cytometry | Thermo Fisher Scientific | Cat# M46753 |
| MitoProbe™ TMRM Assay Kit for Flow Cytometry | Thermo Fisher Scientific | Cat# M20036 |
| FreeStyle 293 Expression Medium | Thermo Fisher Scientific | Cat# 12338018 |
| Benzonase Nuclease | MilliporeSigma | Cat# E1014-25KU |
| PEIpro Transfection Reagent | Polyplus-transfection | Cat# 101000026 |
| UltraPure 1 M Tris-HCl, pH 7.5 | Thermo Fisher Scientific | Cat# 15567027 |
| NaCl (5 M), RNase-free | Thermo Fisher Scientific | Cat# AM9759 |
| UltraPure 0.5 M EDTA, pH 8.0 | Thermo Fisher Scientific | Cat# 15575038 |
| Sucrose, ACS grade | Thermo Fisher Scientific | Cat# 036508.A1 |
| Polyethylene glycol 6,000 | Thermo Fisher Scientific | Cat# A17541.0B |
| Sodium Chloride (Crystalline/Certified ACS) | Fisher | Cat# S271-1 |
| PBS (10X), pH 7.4 | Thermo Fisher Scientific | Cat# 70011044 |
| UltraPure DNase/RNase-Free Distilled Water | Thermo Fisher Scientific | Cat# 10977015 |
| Bolt 4X LDS Sample Buffer | Thermo Fisher Scientific | Cat# B0007 |
| Bolt 10X Sample Reducing Agent | Thermo Fisher Scientific | Cat# B0009 |
| Bolt 20X MES SDS Buffer | Thermo Fisher Scientific | Cat# B0002 |
| Pierce Bovine Serum Albumin Standard Ampules (2 mg/mL) | Thermo Fisher Scientific | Cat# 23209 |
| Protein Standard Mix 15–600 kDa | MilliporeSigma | Cat# 69385 |
| B-PER Bacterial Protein Extraction Reagent | Thermo Fisher Scientific | Cat# 78266 |
| cOmplete EDTA-free Protease Inhibitor Cocktail | MilliporeSigma | Cat# 11836170001 |
| Lysozyme (50 mg/mL) | Thermo Fisher Scientific | Cat# 90082 |
| Benzonase Endonuclease (>90% Purity, HC) | MilliporeSigma | Cat# 71205-3 |
| Carbenicillin Solution (100 mg/mL) | Teknova | Cat# C2136 |
| Chloramphenicol Solution (34 mg/mL) | Teknova | Cat# C0322 |
| 100X Glucose, Glycerol, IPTG Induction Supplement | Teknova | Cat# G0560 |
| Antifoam 204 (10% v/v) | Teknova | Cat# A6427 |
| Blocker BSA (10%) in PBS | Thermo Fisher | Cat #37525 |
| His-Avi-CBLB[39-426] protein | Acro Biosystems | Cat #CBB-H81Q8 |
| Biological Samples |  |  |
| Human Peripheral Blood Leukopak, Fresh | STEMCELL Technologies | Cat# 70500 |
| Experimental models: Cell lines |  |  |
| H1975 | ATCC | Cat# CRL-5908;<br>RRID:CVCL_1511 |
| H1975-mKate2 | ATCC; Modified cells at Outpace Bio | Based on Cat# CRL-5908; RRID:CVCL_1511 |
| H226-A02:01-PRAME+-mKate2 | ATCC; Modified cells at Outpace Bio | Based on Cat# CRL-5826; RRID:CVCL_1544 |
| H1650-mKate2 | ATCC; Modified cells at Outpace Bio | Based on Cat# CRL-5883; RRID:CVCL_1483 |
| Suspension-adapted HEK293T | ATCC; Modified cells at Outpace Bio | Based on Cat# CRL-3216 |
| Experimental models: Organisms/strains |  |  |
| BL21(DE3) | Thermo Fisher Scientific | Cat# C600003 |
| CVB-T7 POL | Avidity | Cat# CVB-T7POL |
| *Saccharomyces cerevisiae* | ATCC | Cat# MYA-4941 |
| NOD.Cg-PrkdcscidIl2rgtm1Wjl/SzJ (NSG) mice | Jackson Laboratory | RRID:IMSR_JAX:005557 |
| NOD-Prkdcem26Cd52Il2rgem26Cd22/NjuCrI (NCG) mice | Charles River Laboratories | RRID:IMSR_CRL:572 |
| Software and algorithms |  |  |
| Prism Version 9 | Graphpad | N/A |
| FlowJo V10.7.1 | BD Biosciences | N/A |
| Everest (Ze5 Cell Analyzer) | Bio-rad | N/A |
| Attune Cytometric Software v6.2.2 (Attune NxT) | Thermo Fisher Scientific | N/A |
| Incucyte 2024B (IncuCyte) | Sartorius | N/A |
| Methodical Mind | Meso Scale Discovery | N/A |
| Octet | Sartorius Octet Analysis Studio v12.2.2.26 | N/A |
| Adobe Illustrator | Abobe | N/A |
| RFDiffusion | RosettaCommons | N/A |
| ProteinMPNN | https://github.com/dauparas/ProteinMPNN | N/A |
| Rosetta | RosettaCommons | N/A |
| RIFDock | https://github.com/rifdock/rifdock | N/A |
| PatchDock | Schneidman Lab | N/A |
| AlphaFold2 | Google DeepMind | N/A |
| PyMOL Molecular Graphics System, Version 1.2r3pre | Schrödinger, LLC. | N/A |
| Other |  |  |
| PrimeSurface 96U 3D Cell Culture Plate | SBio | Cat# MS-9096UZ |
| Streptavidin (SA) Biosensors | Sartorius | Cat# 18-5019 |
| High Precision Streptavidin (SAX) Biosensors | Sartorius | Cat# 18-5117 |
| High Precision Streptavidin 2.0 (SAX2) Biosensors | Sartorius | Cat# 18-5136 |
| Amicon Pro Purification System | MilliporeSigma | Cat# ACS500024 |
| XBridge Premier Protein SEC Column, 250 Å, 2.5 µm, 7.8 x 300 mm | Waters | Cat# 186009962 |
| MaxPeak Premier Protein SEC Guard Column, 250 Å, 2.5 µm, 4.6 x 30 mm | Waters | Cat# 186009969 |
| Superdex 200 Increase 10/300 GL Column | Cytiva | Cat# 28990944 |
| HiLoad 16/600 Superdex 75 pg Column | Cytiva | Cat# 28989333 |
| HisTrap FF 1 mL Column | Cytiva | Cat# 17531901 |
| HisTrap FF 5 mL Column | Cytiva | Cat# 17525501 |

### EXPERIMENTAL MODEL AND SUBJECT DETAILS

#### Animals

NOD.Cg-*Prkdc^scid^Il2rg^tm1Wjl^*/SzJ (NSG) mice (Jackson Laboratory: 005557) NOD-*Prkdc^em26Cd52^Il2rg^em26Cd^*^22^/NjuCrl (NCG) mice (Charles River: 572) All mice were naïve female between 8 and 12 weeks of age at start of experimentation. All experiments were performed under an externally reviewed and approved IACUC protocol at an AAALAC accredited vivarium and in accordance with all local, state, and federal regulations.

#### Cell lines, microbial strains, and culture maintenance

*Adherent Human Tumor Cell Lines*

Human non-small cell lung cancer (NSCLC) cell lines NCI-H1975 (ATCC, CRL-5908), NCI-H1650 (ATCC, CRL-5883), and NCI-H226 (ATCC, CRL-5826) were purchased from the American Type Culture Collection (ATCC) and maintained in complete R10 media comprising RPMI 1640 Medium with GlutaMAX Supplement (Thermo Fisher Scientific) supplemented with 10% v/v certified heat-inactivated Fetal Bovine Serum (FBS, US origin; Thermo Fisher Scientific). Cultures were maintained in tissue culture-treated flasks at 37℃ in a humidified atmosphere containing 5% CO_2_. Cells were routinely passaged at 80-90% confluence using TrypLE Express Enzyme (1x, phenol red; Thermo Fisher Scientific).

For *in vitro* live-cell imaging and cytotoxicity assays (IncuCyte), NCI-H1975 (ROR1+), NCI-H1650 (PRAME+), and NCI-H226 (PRAME-) cells were stably transduced with a lentiviral vector encoding mKate2 fluorescent reporter under the control of an EF1ɑ promoter to generate H1975-mKate2 and H1650-mKate2 target lines. Transduced cells were selected with puromycin dihydrochloride (1.5 ug/mL; Thermo Fisher Scientific) for 10-14 days. mKate2 expression in transduced, antibiotic-selected cells was validated by flow cytometry (>95% mKate2+), then cells were cryopreserved for later use.

For *in vivo* xenograft models evaluating PRAME-specific TCR-T cell efficacy, H226-mKate2-HLA*02:01-PRAME+ cells were generated by first transducing previously generated H226-mKate2 cells with lentivirus encoding human HLA-A*02:01 at a multiplicity of infection (MOI) of one. Surface expression of HLA-A*02:01 was evaluated 72 hours post-transduction by flow cytometry (anti-HLA-A2-BV421, Fixable live/dead eFluor780). Cultures were then subjected to Hygromycin B antibiotic selection (10 ug/mL) until achieving >95% HLA-A2+ purity. The resulting NCI-H226.HLA-A*02:01 intermediate line was subsequently transduced (MOI = 1) with lentivirus encoding full-length human PRAME. At day 3 post-transduction, cells were selected with Blasticidin (10 ug/mL) for 2 weeks. Ectopic PRAME expression in selected bulk populations was verified by isolating total mRNA and quantifying transcripts via reverse transcription-digital droplet PCR (RT-dPCR). H226-PRAME+-HLA-A*02:01-mKate2 purified cells were then cryopreserved for later use. For in vivo ROR1 CAR-T cell xenograft evaluation, unmodified parental H1975 (ROR1+) cells were utilized. After thawing cells for use in in vivo xenograft models, all unmodified as well as engineered tumor cell lines were maintained under standard R10 culture conditions (no antibiotics) and harvested during exponential growth phase (Passages 3–8) prior to subcutaneous inoculation.

#### Suspension-Adapted HEK293T Line for Lentiviral Production

Suspension-adapted HEK293T cells utilized for high-titer lentiviral production were cultured in FreeStyle 293 Expression Medium (Thermo Fisher Scientific) in polycarbonate vented-cap Erlenmeyer flasks (Corning). Cultures were maintained on an orbital shaker (130 rpm, 50 mm orbit stroke) at 37℃ in 8% CO_2_ and 80% relative humidity. Cells were routinely passaged every 3-4 days at a density of 0.2-0.4 x 10^6 cells/mL and maintained in exponential growth phase (>90% viability) prior to transfection.

#### Yeast Surface Display Strain (Saccharomyces cerevisiae)

Saccharomyces cerevisiae EBY100 MATa AGA1::GAL1-AGA1::URA3 ura3-52 trp1 leu2-delta1 his3-delta200 pep4::HIS3 prbd1.6R can1 GAL (ATCC) was utilized for yeast surface display library screening and binder selections. EBY100 cells were routinely maintained on YPD agar plates (1% w/v yeast extract, 2% w/v peptone, 2% w/v dextrose, 2% w/v agar) at 30℃. Transformed yeast display libraries were grown in synthetic dextrose minimal medium with casamino acids (SD-CAA; 2% w/v dextrose, 0.67% w/v yeast nitrogen base without amino acids, 0.5% w/v casamino acids, 0.54% w/v Na_2_HPO_4_, 0.86% w/v NaH_2_PO_4_·H_2_O) at 30℃ with shaking (250 rpm). Cell-surface expression of display constructs was induced by transferring exponentially growing cells into synthetic galactose minimal medium with casamino acids (SG-CAA; identical to SD-CAA except substituting 2% w/v galactose for dextrose) at an initial OD600 of 0.5-1.0, followed by incubation at 30℃ for 18-24 hours.

#### Recombinant Protein Expression Strain (Escherichia coli)

BL21(DE3) were transformed with plasmids encoding the protein of interest by heat shock and recovered in SOC medium at 37℃ prior to culture in LB medium supplemented with 100µg/mL carbenicillin and grown for approximately 16 hours. Expression cultures were inoculated at a volumetric ratio of 1:60 confluent culture:Autoinduction media. Autoinduction media was made with either ZY-5052 Autoinduction Media Base (Teknova) and 120mL/L of 3S8600 Autoinduction Supplement (Teknova) or Terrific Broth and 10mL/L of G0560 Induction Supplement (Teknova). Autoinduction media was also supplemented with 100µg/mL carbenicillin, 0.005% Anti-Foam 204, and 50µM Biotin. Cultures were grown at 37℃ for 4 hours with shaking at 300-350 rpm and shifted to 18℃ for 20-24 hours more before harvesting by centrifugation.

CVB-T7-POL were transformed with plasmids encoding the protein of interest by heat shock and recovered in SOC medium at 37℃ prior to culture in LB medium supplemented with 100µg/mL carbenicillin and grown for approximately 16 hours. Expression cultures were inoculated at a volumetric ratio of 1:60 confluent culture:Autoinduction media. Autoinduction media was made with either ZY-5052 Autoinduction Media Base (Teknova) and 120mL/L of 3S8600 Autoinduction Supplement (Teknova) or Terrific Broth and 10mL/L of G0560 Induction Supplement (Teknova). Autoinduction media was also supplemented with 100µg/mL carbenicillin,10µg/mL chloramphenicol, 0.005% Anti-Foam 204 and 50µM Biotin. Cultures were grown at 37℃ for 4 hours with shaking at 300-350 rpm and shifted to 18℃ for 20-24 hours more before harvesting by centrifugation.

#### Primary human T cell isolation and culture

Normal donor leukopaks (StemCell) were processed within 48h of receipt. Apheresis bags were mixed thoroughly and transferred to an appropriately sized conical. Cells were resuspended at a concentration of 50E6 to 75E6 cells/mL with EasySep Buffer (StemCell). A portion of the suspension was banked directly as frozen PBMCs. Isolated T cells used in downstream assays showed >95% CD3^+^.

### METHOD DETAILS

#### Computational design of pan-NR4A *de novo* protein binders

Structural motifs that are conserved across the ligand-binding domain (LBD) of NR4A1, NR4A2, and NR4A3 were identified (FIG. 2). Binder backbone structures were generated using RFDiffusion^18^. PDB ID 3v3e was used as the targeting structure and residues 574,575,592,593,596,598 (UNIPROT P22736 numbering) near the C-terminus were selected as hotspot residues. 10,000 diffusion trajectories designing 80 amino-acid binders were produced and the structures from several early timesteps were output in addition to the final structure. The amino acid sidechains of these outputs were designed with ProteinMPNN^19^, filtered by Rosetta ddG, and 300,000 were predicted with AlphaFold2 initial guess^21^. The best 300 structures (by AF2 pae_interaction) had their interface helices extracted and a further 10,000 diffusion trajectories (spread evenly across the 300) were used to design new binders around these interface helices ranging from 65aa-120aa. These outputs were designed with ProteinMPNN, filtered by Rosetta ddG, and 300,000 were predicted with AlphaFold2 initial guess. Pooling the 600,000 designs together, the best 7,500 were selected by a combination of Rosetta ddG and AlphaFold2 pae_interaction and were tested for binding against purified recombinant LBD of NR4A1 and NR4A3 by yeast surface display.

#### Computational design of *de novo* Cbl protein binders

Designs were generated using the PatchDock + RifDock protocol from Cao et. al 2022 [https://www.nature.com/articles/s41586-022-04654-9]. Three sites were selected:

|  | PDB | Patchdock Res (Uniprot) | Rifdock Res (uniprot) |
| --- | --- | --- | --- |
| Site1 | 3PFV | 109,192,193,196,227 | 105,108,109,191,192,193,196,226,227 |
| Site2 | 3ZNI | 50,57,75,79,82 | 50,53,54,57,61,75,79,82,83 |
| Site3 | 3PFV | 181,185,204,243 | 177,180,181,184,185,203,204,205,243 |

A library of 5000 50-60aa 3-helical bundles was docked with patchdock using the Patchdock Res above to generate around 1000 docks per scaffold. The docks were passed into RifDock for refinement and around 5,000,000 total docks per site were output. These were passed through a rough Rosetta filter looking for hydrophobic contacts with the PatchDock res to cut the number down to 500,000 which were run through Rosetta FastDesign. These 500,000 were subsequently filtered by ddG as well as contact_molecular_surface to the hydrophobic contacts with the best 200,000 being filtered by AF2 with initial guess (Nate Bennett Nature 2023). The best 3000 motifs were selected by a combination of AF2 pae_interaction and ddG. These 3,000 motifs were combined with the 5000 scaffolds to generate around 5,000,000 new docks. These new docks were then roughly filtered to 500,000, FastDesigned, filtered by ddG, and predicted with AF2. The best 15,000 total outputs by AF2 pae_interaction were selected from the combination of the RifDock round and the motif grafting round. Designs were visually inspected using the PyMOL Molecular Graphics System, Version 1.2r3pre, Schrödinger, LLC.

#### Construction of Site Saturation Mutagenesis (SSM) library for Cbl binders

Preliminary yeast display data identified 3 binders to each of site1 and site2 that were predicted to be weak affinity. A subsequent SSM library was constructed making every mutation at every position for each of the 6 proteins. Before constructing the SSM library, a polishing step whereby AF2 and Rosetta were used to make mutations in a monte-carlo fashion were employed to generate further mutants to test for the SSM library. Upon sorting this library, it was seen that 4 of the 6 binders were very weak, leaving 1 site 1 and 1 site 2 binders of which only the site 2 passed rigorous numerical SSM validation tests where Rosetta’s ability to predict the effect of each mutation is compared to experimental data. (see Cao et. al 2022) The sequence polished version of the best binder was used as the basis for the next step. The SSM revealed many mutations that were predicted to improve binding. We selected the mutations 2IN, 7HEDQ, 8NTAHPD, 12SA, 13L*EV, 15HD, 16GE, 17LV, 18ST, 19GA, 21GA, 23HQ, 27KR, 31LH, 51IR to form the basis of the combo library. The library was sorted for 3 rounds collecting only the highest-affinity binders. Sanger sequencing was used to identify the top hits which were subsequently tested for binding kinetics by BLI.

#### Yeast display

Chip-synthesized oligonucleotides ordered from Agilent Technologies were used to amplify the designed DNA fragments. For designs with sequences longer than, two-fragment assembly was used to assemble the genes using the method described in Basanta et. al 2020 (https://www.pnas.org/doi/10.1073/pnas.2005412117#sec-3). The library DNA was combined with linearized pETCON3 vector and transformed into an EBY100 yeast strain by electroporation. Using the Sony SH800S instrument, the library was first sorted for expression by labeling cells with anti-c-Myc-FITC conjugated antibody (Immunology Consultants Laboratory) and collecting the desired population. Subsequent sorts were performed by labeling cells for expression and binding in the presence of biotinylated target protein, anti-c-Myc-FITC conjugated antibody, and Streptavidin R-Phycoerythrin (SAPE, Thermo Fisher Scientific). For NR4A, cells from the expression sort were induced and labeled at 1000nM without avidity with the target protein in PBSF (phosphate buffered saline with 1 % (w/v) BSA) for round one, collecting yeast cells that displayed positive signals for both anti-c-Myc-FITC and SAPE. The second round of sorting was performed at various concentrations (titration) decreasing 10-fold from 1000nM to 0.1nM. For Cbl-b, cells from the expression sort were induced and labeled at 1000nM with avidity for the initial two rounds of sorting. The third round of sorting was a titration performed at various concentrations decreasing ∼3-fold from 3000nM to 400nM. To generate the SSM library for Cbl-b, oligonucleotides were ordered from Agilent Technologies. DNA was amplified and the yeast display library sorted as previously described. In brief, the library was first sorted for expression, followed by sort one at 1000nM without avidity, and a titration sort decreasing 5-fold from 1000nM to 0.2nM for round two. All collected library samples were prepared and sequenced using Illumina Nextseq or MiSeq sequencing. Using the sequencing data gathered from yeast display, SC_50_ values were estimated for each binder in the library as previously described in Cao et. al 2022 [https://www.nature.com/articles/s41586-022-04654-9]

Note that for NR4A binders, no site-saturation-mutagenesis (SSM) or combination libraries were required; monomeric NR4A target protein was used for yeast display and sorting (avid-tetramerized NR4A target protein was poorly behaved).

#### Generation of an NR4A3-specific monobody using phage display and yeast affinity maturation

A 2.95 × 10^10^ member FN3-based (uniprot ID P02751 aa1538-1631 D1540S) synthetic monobody library with NNK-randomized BC, DE, and FG loops was panned against NR4A3-LBD for four rounds. Polyclonal phagemid DNA from R4 was amplified by error-prone PCR (GeneMorph II, Agilent), shuffled, and the resulting library was cloned into the pETCONV3 vector, transformed into EBY100, and sorted against NR4A3-LBD for four rounds. Following sorting, Sanger sequencing was used to identify molecules for kinetic and biophysical characterization.

#### Recombinant protein expression & purification

Truncated NR4A1[361-598], NR4A2[363-598] and NR4A3[395-626] proteins were fused to N-terminal Avi-6His tags. Pan-NR4A de novo binders, nr77 and nr78, and their associated interface mutants, nr77mut and nr78mut, were fused to an N-terminal 6His-TEV tag. Cbl-b de novo binders were fused to an N-term 6His-MBP tag. The NR4A3 monobody was fused to an N-terminal 6His-SUMOStar tag. All were expressed in E coli in BL21(DE3) or CVB-T7-POL cells. Cultures were grown in autoinduction media with expression at 18°C. Biotinylated target proteins were biotinylated in vivo via coexpression of BirA in the presence of 50µM biotin. Harvested cells were lysed either mechanically via microfluidizer (in the presence of protease inhibitors and Benzonase) or chemically using B-PER (Phosphate). Clarified, filtered lysates were subjected to immobilized metal affinity chromatography (IMAC) with Ni-NTA resin in batch or FPLC column format. For The NR4A3 monobody cleavage of the SumoStar and His tag was done overnight at 4°C with SumoStar Protease and a second nickel affinity purification in PBS for removal of tag and protease from protein samples prior to filtration and aliquot preparation. Cbl-b de novo binders were concentrated directly after IMAC. For NR4A de novo binders and NR4A1–3 target proteins, IMAC eluates were further purified by size-exclusion chromatography (SEC) using Superdex 75 or 200 columns equilibrated in target-specific buffers (HEPES with TCEP for NR4A1–3; PBS for NR4ABinders). After purification protein samples were concentrated using 3K or 10K MWCO spin filters, 0.2 μm filtered, aliquoted, and stored at −80°C. Purity was confirmed using SDS-PAGE and HPLC-SEC. Biotinylated human CBL-B was purchased from Acro (CBB-H81Q8).

#### Binding kinetics measurements by Biolayer interferometry (BLI, Octet)

Binding of *de novo* and monobody binders to NR4A1, NR4A2, NR4A3, or Cbl-b, target protein was measured using BLI on an Octet RED96 system (Sartorius). All measurements were carried out at 25°C in a running buffer of 1X PBS pH 7.2 supplemented with 0.5% BSA. Biotinylated target proteins were captured on streptavidin (SA, SAX, or SAX2) biosensors such that theoretical Rmax values would be 0.2-1 nm. Binding of captured target ligands to miniprotein analytes was measured at multiple concentrations for each interaction. Concentrations tested were: 50-1.56 nM (2-fold dilutions), 12-0.375 nM (2-fold dilutions), 1000-0.98 nM (4-fold dilutions), and 1000-0.98 nM (4-fold dilutions) for nr77, nr78, Cbl *de novo* binders, and the NR4A3 monobody, respectively. Interface mutants of nr77 and nr78 were also tested at 50 nM and 12 nM to demonstrate perturbation of binding. [Association and dissociation] times were: [600 sec, 1800 sec], [600 sec, 1800 sec], [240 sec, 600 sec], and [300 sec, 600 sec] for nr77, nr78, Cbl *de novo* binders, and the NR4A3 monobody, respectively. Data were fit globally using a 1:1 kinetic binding model using Sartorius Octet Analysis Studio v12.2.2.26

#### Lentiviral vector production, concentration, and functional titering

Suspension-adapted HEK293T cells were seeded 24 hours prior to transfection in pre-warmed FreeStyle 293 Expression Medium at a density of 1 x 10^6 cells/mL to achieve exponential growth and reach approximately 2 x 10^6 cells/mL at the time of transfection. Cultures were prepared across three operational scales:

● Small-scale (2mL final yield): Cells were seeded in an initial volume of 2mL in 24-well shake plates.
● Medium-scale (120mL final yield): Cells were seeded in an initial volume of 60mL in 500mL Optimum Growth Flasks.
● Large-scale (320mL final yield): Cells were seeded in an initial volume of 160 mL in 1.6 L Optimum Growth Flasks.

At 24 hours post-seeding, cells were transfected via polyethylenimine (PEI)-mediated transient transfection using PEIpro (Polyplus-transfection). Master mixes containing third-generation lentiviral packaging plasmids, transfer plasmid DNA, and ambient-temperature FreeStyle 293 medium were combined with PEIpro diluted in FreeStyle 293 medium. For flask-scale preparations, transfection complexes were mixed by gentle inversion (5–6 times); for 24-well shake plates, the PEI/medium mix was added dropwise to the DNA solution. Transfection complexes were incubated for 10min at room temperature before dropwise addition to cell suspensions. At 18 hours post-transfection, flask cultures were supplemented 1:1 with fresh, pre-warmed FreeStyle 293 medium containing 10 U/mL Benzonase Nuclease (MilliporeSigma; final working concentration 5 U/mL). No additional medium expansion was performed for 2mL shake plate cultures. At 72 hours post-transfection, lentiviral supernatants were harvested and clarified by centrifugation at 300 x g for 10 min at room temperature. Clarification and concentration proceeded according to production volume:

2mL scale: Clarified supernatant was filtered directly using a 0.22 um polyethersulfone (PES) Millex syringe filter into a 15 mL conical tube. Viral particles were concentrated by adding Lentivirus PEG-6000 concentrator at a 3:1 (v/v) ratio, mixed by inversion, and incubated at 4℃ for 2 hours. Samples were centrifuged at 3,000 x g for 1.5 hours at 4℃. Supernatants were carefully aspirated, and viral pellets were resuspended in cytokine-free T cell media to achieve a 10x concentration relative to the harvest volume.

Flask scale (120mL or 320mL final yield): Clarified supernatants were vacuum-filtered using 0.22 um PES Nalgene Rapid-Flow Sterile Disposable Filter Units (500mL or 1000mL capacity, Thermo Scientific) and pooled. Supernatants were underlaid with a 10% (w/v) sucrose cushion at a 4:1 (v/v) ratio in 50 mL conical tubes and concentrated by ultracentrifugation at 10,000 x g for 4 hours at 4℃. Supernatants were decanted, and pellets were resuspended in Final Formulation Buffer (cytokine-free T cell media supplemented with 5% Lentivirus PEG-6000 concentrator) to yield a 100x concentration. Resuspended pellets were vortexed vigorously and incubated overnight at 4℃ to ensure full dissolution.

All concentrated viral batches were pooled, aliquoted, and stored at −80℃. Functional viral titers were determined by transducing Jurkat T cells or primary human T cells at limiting dilutions. Expression of the transduction marker EGFR, CD19, or Vb8 was quantified by flow cytometry at day 2 (Jurkat) or day 4 (primary T cells) post-transduction. Functional transducing units per milliliter (TU/mL) were calculated from transduction marker expression (% positive cells) according to the following formula:

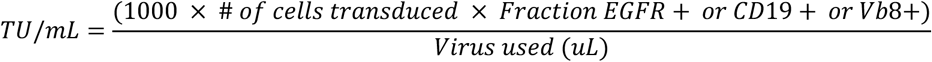

#### Flow-based degradation assay of exogenous NR4A1-HA, NR4A2-HA, and NR4A3-HA

##### Cell Thawing and Activation

Cryopreserved primary T cells were thawed in a 37°C water bath then diluted 1:10 in T cell media (OpTmizer T Cell Expansion SFM supplemented with Immune Cell Serum Replacement, 2 nM Glutamine, and 1x GlutaMAX). Cells were then centrifuged at 300 x g for 10 minutes, and pellets were resuspended and counted using an automated cell counter. Cell suspensions were then adjusted to 2 x 10^6 live cells/mL in T cell media supplemented with recombinant human IL-2 (2 x 10^2 IU/mL), IL-7 (1.2 x 10^3 IU/mL), IL-15 (2 x 10^2 IU/mL), and TransAct (10 µL/mL). Cell suspensions were transferred to a culture flask and incubated at 37°C in 5% CO_2_ overnight.

##### Dual Lentiviral Transduction

Following activation, T cells were centrifuged at 300 x g for 10 minutes and adjusted to 1 x 10^5 live cells per 100 µL in cytokine-supplemented media containing TransAct. Cells were aliquoted into microplates, and concentrated lentivirus encoding the NR4A-HA targets (CD19+ or EGFR+) was added (MOI=10). After a 4–5 hour incubation at 37°C in 5% CO_2_, 100 µL of the cell suspension was transferred to a 96-well flat-bottom plate, and either 100 µL of unconcentrated lentivirus (DHD37 heterodimer degrader library screening) or 10 µL of concentrated lentivirus (nr77 vs. nr78 binder-degrader comparison) encoding binder-degrader proteins (CD19+ or EGFR+, respectively) was added. Dual-transduced cells were incubated overnight at 37°C in 5% CO_2_.

##### Expansion, Maintenance, and MLN4924 Treatment

To mitigate TransAct-mediated toxicity, 100 µL of the cell suspension was transferred to a new 96-well plate and diluted with 100 µL of fresh cytokine-supplemented T cell media. For DHD37B heterodimer degrader library screening: Cells were incubated for two days at 37°C in 5% CO_2_. For nr77 vs. nr78 binder-degrader comparison: Cells were incubated for one day at 37°C in 5% CO_2_, then the following day 100 µL of cells were transferred to a new 96 well flat bottom plate. 100 µL of cytokine-supplemented media was added to one plate, and as an additional control 100 µL of cytokine-supplemented media containing MLN4924 (0.2uM), a NEDDylation inhibitor that prevents the activation of Cullin-RING ligases (CRLs), was added to the duplicated plate to block SPOP-mediated degradation. Cells were then incubated for an additional day at 37°C in 5% CO_2_.

##### Flow Cytometry Staining and Data Acquisition

Three days post-transduction, cells were harvested and transferred into a 96-well U-bottom plate. Cells were washed twice with BioLegend Cell Staining Buffer (CSB). Surface staining was performed by incubating cells with a cocktail containing a fixable viability dye (FVD780), anti-CD19-FITC, and anti-EGFR-BV421 antibodies in CSB for 20–30 minutes at room temperature in the dark.

Following surface staining, cells were washed twice in CSB and fixed/permeabilized using the Foxp3/Transcription Factor Staining Buffer Set (Thermo Fisher Scientific) according to the manufacturer’s protocol. Intracellular staining for the HA and FLAG tags was performed by incubating cells with anti-HA-AF647 and anti-FLAG-PE antibodies in 1x Perm/Wash buffer for 45 minutes at 4°C in the dark. Cells were washed twice in 1x Perm/Wash buffer and resuspended in CSB for acquisition. Data were collected using an Attune NxT Flow Cytometer (Thermo Fisher Scientific) equipped with an autosampler.

##### DHD37 heterodimer degrader library screening data analysis

Data analysis was performed using FlowJo software. To quantify NR4A-HA protein degradation, the Median Fluorescence Intensity (MFI) of the HA tag was calculated across two distinct gated populations: dual-transduced cells (CD19+/EGFR+), and non-transduced internal controls (CD19-). A Targeted Degradation Score (TDS) was calculated to represent a ratio of the background-subtracted HA MFI in DHD37B-degrader expressing cells relative to a DHD37B without degrader (non-degrading) expressing control:

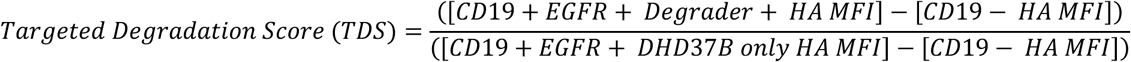

The CD19-population served as the internal control for background fluorescence.

##### nr77 vs. nr78 binder-degrader comparison data analysis

Data analysis was performed using FlowJo software. To quantify NR4A-HA protein degradation, the Median Fluorescence Intensity (MFI) of the HA tag was calculated across three distinct gated populations: dual-transduced cells (CD19+/EGFR+), target-only expressing cells (CD19-/EGFR+), and target protein non-transduced internal controls (EGFR-). A Targeted Degradation Score (TDS) was calculated to represent the ratio of background-subtracted HA MFI in degrader-expressing cells relative to the target-only expressing control:

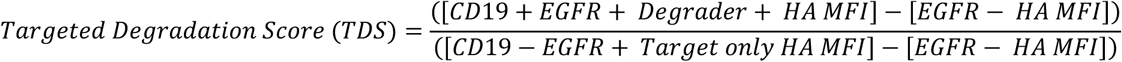

The EGFR-population served as the internal control for background fluorescence.

#### Flow-based assay to assess degradation of endogenous NR4A1 in primary human T cells

##### Cell Thawing and Activation

Cryopreserved primary T cells were thawed in a 37°C water bath, diluted 1:10 in T cell media, then centrifuged at 300 x g for 10 minutes. Pellets were resuspended and counted using an automated cell counter. Cell suspensions were then adjusted to 2 x 10^6 live cells/mL in T cell media supplemented with recombinant human IL-2, IL-7, IL-15, and TransAct, then incubated at 37°C in 5% CO_2_ overnight.

##### Lentiviral Transduction

Following activation, T cells were centrifuged at 300 x g for 10 minutes and adjusted to 1 x 10^5 live cells per 100 µL in cytokine-supplemented media containing TransAct. 100 µL of the cell suspension was transferred to a 96-well flat-bottom plate, and 10 µL of concentrated lentivirus encoding binder-degrader proteins (CD19+) was added. Transduced cells were incubated overnight at 37°C in 5% CO_2_.

##### Expansion, Maintenance, and MLN4924 Treatment

To mitigate TransAct-mediated toxicity, 60 µL of cells were transferred to a new 96-well flat bottom plate and diluted with 140 µL of fresh cytokine-supplemented T cell media. Cells were then incubated for three days at 37°C in 5% CO_2_. Following expansion, 100 µL of cells were transferred to a new 96 well flat bottom plate, diluted with 100 µL of fresh cytokine-supplemented T cell media, then incubated for an additional two days at 37°C in 5% CO_2_. After two days of additional expansion, 100 µL of cells were transferred to a new 96 well flat bottom plate. 100 µL of cytokine-supplemented media was added to one plate, and as an additional control 100 µL of cytokine-supplemented media containing MLN4924 (0.2uM final concentration), a NEDDylation inhibitor that prevents the activation of Cullin-RING ligases (CRLs), was added to the duplicated plate to block SPOP-mediated degradation. Cells were then incubated for an additional day at 37°C in 5% CO_2_.

##### Induction of Endogenous NR4A1, Flow Cytometry Staining and Data Acquisition

7 days post-transduction, a pooled sample from each sample group with or without MLN4924 treatment was generated to be used as an untreated control (No endogenous NR4A1 upregulation). To all other cells 100 µL of TransAct supplemented T cell media was added and cells were incubated for 3 hours at 37°C in 5% CO_2_ in order to induce endogenous NR4A1 upregulation. After TransAct treatment cells were harvested and transferred into a 96-well U-bottom plate. Cells were washed twice with BioLegend Cell Staining Buffer (CSB). Surface staining was performed by incubating cells with a cocktail containing a fixable viability dye (FVD780) and anti-CD19-FITC antibodies in CSB for 20–30 minutes at room temperature in the dark.

Following surface staining, cells were washed twice in CSB and fixed/permeabilized using the Foxp3/Transcription Factor Staining Buffer Set (Thermo Fisher Scientific) according to the manufacturer’s protocol. Intracellular staining for endogenous NR4A1 was performed by incubating cells with anti-NR4A1-PE antibody in 1x Perm/Wash buffer for 45 minutes at 4°C in the dark.

Cells were washed twice in 1x Perm/Wash buffer and resuspended in CSB for acquisition. Data were collected using an Attune NxT Flow Cytometer (Thermo Fisher Scientific) equipped with an autosampler.

#### Data Analysis

Data analysis was performed using FlowJo software. To quantify the degradation of endogenous NR4A1, the Median Fluorescence Intensity (MFI) of NR4A1-PE was calculated across three distinct gated populations: TransAct-treated, binder-degrader expressing (CD19+) cells, TransAct-treated, non-transduced (CD19-) internal positive controls, and untreated, pooled (CD19-) negative controls. A Targeted Degradation Score (TDS) was calculated as the ratio of background-subtracted NR4A1 MFI in degrader-expressing cells relative to the non-degrading (CD19-) internal control:

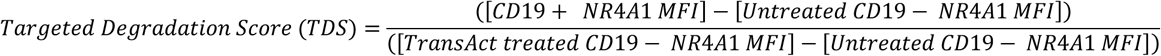

The untreated, pooled (CD19-) cell population served as the internal control for background fluorescence.

##### Flow cytometry for primary T cell production and in vitro chronic stimulation assays

Cell samples were transferred to 96-well U-bottom plates and washed once with 150 uL Cell Staining Buffer (CSB; BioLegend) by centrifugation at 500 x g for 2 min at room temperature.

For antigen-specific T cell identification using MHC dextramer, cells were first resuspended in 30 uL of PRAME-specific Dextramer-PE (2 uL stock per 50 uL total volume in CSB; Immudex) and incubated for 10 min at room temperature in the dark. Without washing, 20 uL of a 2.5x fluorophore-conjugated surface antibody and fixable live/dead eFluor780 (0.0125-0.025 uL stock per 50 uL total volume; Thermo Fisher Scientific) cocktail was added directly to the dextramer reaction (yielding a final staining volume of 50 uL diluted in CSB), and cells were incubated for an additional 20 min at room temperature in the dark.

For standard panels lacking dextramer staining, cells were resuspended directly in 50 uL of surface antibody and fixable live/dead eFluor780 cocktail prepared in CSB (total staining volume of 50 uL) and incubated for 20-30 min at room temperature in the dark.

Following surface staining, cells were washed twice with 150 uL CSB, resuspended in FluoroFix Buffer (BioLegend), and incubated at 4℃ for at least 30 min to fix the cells. Flow cytometric acquisition was performed on an Attune NxT Flow Cytometer (Thermo Fisher Scientific) or a ZE5 Cell Analyzer (Bio-Rad). All flow cytometry data were analyzed using FlowJo software (BD Biosciences).

Antibody dilutions used for flow staining that aren’t described explicitly in the Method Details section are outlined in the Key Resources Table (KRT).

##### Subsequent lentiviral transduction and dual bead purification of primary T cells

*Cell Thawing and Activation*

Cryopreserved primary T cells were thawed in a 37°C water bath, diluted 1:10 in T cell media, then centrifuged at 300 x g for 10 minutes. Pellets were resuspended and counted using an automated cell counter. Cell suspensions were then adjusted to 2 x 10^6 live cells/mL in T cell media supplemented with human IL-2, IL-7, IL-15, and TransAct, then incubated at 37°C in 5% CO_2_ overnight. Cells were then transduced and prepared as follows:

##### Ror1 CAR-tEGFR Lentiviral Transduction

After overnight activation cell suspensions were centrifuged at 300xg for 10 minutes, and pellets were resuspended in T cell media then counted using an automated cell counter. Cell suspensions were then adjusted to the desired live cell concentration in T cell media supplemented with human IL-2, IL-7, IL-15, and TransAct. Cells were then aliquoted into a microwell plate and subsequently transduced with a Ror1 CAR-tEGFR construct (MOI=10), then expanded for 4 days.

##### anti-EGFR bead purification

After expansion for 4 days Ror1 CAR transduced cells were anti-EGFR bead-purified to select for Ror1 CAR+ cells as follows: Cells were centrifuged at 300 x g for 10 min at 4°C, then cell pellets were resuspended in cold MojoSort Buffer (BioLegend) supplemented with a biotinylated anti-human EGFR antibody (BioLegend) at a final concentration of 1:500 (200uL of antibody solution per 10 x 10^6 live cells). Cells were incubated on ice for 30 min. Following primary incubation, 2mL of cold MojoSort Buffer was added, and cells were centrifuged at 1,500 x g for 1 min. Cells pellets were resuspended in cold MojoSort Buffer (100uL/1×10^6 cells) containing MojoSort Streptavidin Nanobeads (BioLegend) at 8uL Nanobeads/100uL buffer. Cell-bead suspensions were incubated on ice for 15 min, then washed twice in 2mL of cold MojoSort Buffer before being transferred to 5 mL round-bottom flow cytometry tubes. For magnetic separation, tubes were placed in an EasySep Magnet (Stemcell Technologies) for 10 min at room temperature. Supernatant containing unbound cells was decanted, then tubes were removed from the magnet and remaining cells were resuspended in 2mL of cold MojoSort Buffer. This magnetic separation cycle was repeated for a total of three sequential rounds to ensure maximum yield and purity of the EGFR+ population. Following the final magnetic selection step, purified EGFR+ Ror1 CAR-T cells were resuspended in 3mL of cold T cell media. Cells were centrifuged at 300 x g for 10 min, supernatant decanted, and pellet resuspended in T cell media.

##### OUTLAST regulator-tCD19 lentiviral transduction and anti-CD19 bead purification

Immediately after anti-EGFR bead purification cells were transduced with OUTLAST regulator-tCD19 constructs (MOI=10) in T cell media supplemented with human IL-2, IL-7, and IL-15. Cells were then expanded for an additional 3 days followed by anti-CD19 bead purification (EasySep Human CD19 Positive Selection Kit II; Stemcell Technologies) according to the manufacturer’s instructions to select for a pure population of dual transduced cells. Cells were then expanded in T cell media supplemented with human IL-2, IL-7, and IL-15 for an additional 3-5 days before proceeding with functional assays using freshly prepared cells. During this expansion the success of bead purification processes was determined by flow cytometry (Fixable live/dead eFluor 780, human Ror1-Fc-AF647, anti-EGFR-BV421, anti-CD19-FITC).

##### All-in-one vector lentiviral transduction of primary T cells

*Cell Thawing and Activation*

Cryopreserved primary T cells were thawed in a 37°C water bath, diluted 1:10 in T cell media, then centrifuged at 300 x g for 10 minutes. Pellets were resuspended and counted using an automated cell counter. Cell suspensions were then adjusted to 2 x 10^6 live cells/mL in T cell media supplemented with recombinant human IL-2, IL-7, IL-15, and TransAct, then incubated at 37°C in 5% CO_2_ overnight. Cells were then transduced and prepared as follows:

*Lentiviral Transduction and Culture Expansion*

After overnight activation cell suspensions were centrifuged at 300xg for 10 minutes, and pellets were resuspended in T cell media then counted using an automated cell counter. Cell suspensions were then adjusted to the desired live cell concentration in T cell media supplemented with human IL-2, IL-7, IL-15, and TransAct, aliquoted into a GREX 24-well plate (Wilson Wolf), and subsequently transduced with all-in-one Ror1 CAR-OUTLAST regulator-tEGFR or PRAME-specific TCR-OUTLAST regulator-tEGFR lentivirus (MOI=10). Transduced T cells were then expanded for 7 days in T cell media supplemented with human IL-2, IL-7, and IL-15, with additional cytokines added on days 3 and 5.

*Sample Normalization and Cryopreservation*

On day 7 transduction efficiency was determined by flow cytometry for Ror1 CAR+EGFR+ (Fixable live/dead eFluor 780, human Ror1-Fc-AF647, anti-EGFR-BV421) or PRAME-specific Dextramer+ (PRAME-specific Dextramer-PE, anti-CD8-BUV395, anti-CD4-BUV496, anti-EGFR-BV421, anti-Vb8-APC, Fixable live/dead eFluor 780, Brilliant Stain Buffer Plus), then all samples were normalized to equivalent Ror1 CAR+EGFR+ or PRAME-specific Dextramer+ frequencies by dilution with donor-matched, untransduced Mock cells. Cells were then centrifuged at 300 x g for 10 minutes, and pellets were reususpended in CryoStor CS10. 1mL aliquots were transferred into cryovials and placed into a CoolCell. CoolCells were transferred to a −80℃ freezer overnight, then transferred to an LN2 tank the following day for long term storage until ready to use in T cell functional assays.

##### Generation and validation of knockout (KO) ROR1 CAR-T Cells

*CRISPR/Cas9 ribonucleoprotein (RNP) assembly*

Lyophilized synthetic sgRNAs (Synthego) were reconstituted per the manufacturer’s instructions and RNPs were assembled by combining recombinant Cas9 protein (Thermo Fisher Scientific) with either multiple or one sgRNA(s) at a Cas9:total-sgRNA molar ratio of approximately 1:8.2. When multiple sgRNAs were used each of the guides contributed an equal share of the total sgRNA. Assembled RNPs were incubated at room temperature for 10–20 minutes prior to electroporation.

##### T Cell Electroporation

On the day of electroporation, a subset of enriched Ror1 CAR-T cells were collected, gently dissociated, counted, and centrifuged (300 × g, 5 min), followed by a single wash in PBS. Cells were resuspended in a proprietary nucleofection buffer (Lonza P3 Primary Cell solution with supplement) at a density of approximately 1 × 10⁶ cells per 20 µL for small-scale reactions. Cell suspensions were combined with the assembled RNP complexes in electroporation cuvettes and electroporated using a 4D-Nucleofector (Lonza) with a pulse code optimized for primary human T cells (DN-100). Following electroporation, cells were diluted in the cuvette with T cell media before being transferred into T cell media supplemented with IL-2, IL-7, and IL-15 in tissue-culture plates and returned to standard culture conditions (37 °C, 5% CO₂). Fresh cytokine-supplemented media was added approximately 24 hours post-electroporation to support recovery and expansion.

##### Genomic DNA extraction

Approximately 48–72 hours after electroporation, an aliquot of cells (2-4 × 10^5^ cells per condition) was collected from each edited population for genotyping, with the remainder retained for downstream functional assays. Genomic DNA was extracted using a column-based genomic DNA purification kit (Zymo Quick-DNA Miniprep) according to the manufacturer’s protocol. DNA concentration and purity were assessed by UV-Vis spectrophotometry.

##### PCR amplification and Sanger sequencing

The genomic region flanking the sgRNA target sites was amplified by PCR from both edited and matched wild-type genomic DNA using a high-fidelity DNA polymerase master mix (Q5 High-Fidelity 2X Master Mix, NEB) with locus-specific forward and reverse primers, following standard thermocycling conditions (initial denaturation at 98 °C; 25–35 cycles of denaturation, primer-specific annealing, and extension; final extension at 72 °C). PCR products were purified using a column-based PCR clean-up kit (Zymo Clean & Concentrator) according to the manufacturer’s protocol and submitted for Sanger sequencing (Genewiz) using a locus-specific sequencing primer positioned downstream of the edited region.

##### Indel quantification and KO scoring

Sanger sequencing traces from edited samples were compared against traces from matched wild-type (unedited) controls from the same donor using Synthego’s Inference of CRISPR Edits (ICE) analysis tool to quantify insertion/deletion (indel) frequencies and estimate the proportion of cells carrying frameshift (KO) alleles at the target locus for each donor.

##### *In vitro* sequential tumor challenge assay

To evaluate long-term functional persistence and serial cytotoxicity, Ror1 CAR-T cells were subjected to repeated tumor antigen challenges using a live-cell imaging assay. On the day of initial assay setup, H1975-mKate2 target cells were seeded onto 96 well flat bottom tissue culture plates at 2 x 10^4 cells/well in 100uL R10 media and incubated for 2–3 hours to permit cell adherence. Freshly transduced and bead-purified primary T cells expressing the ROR1 CAR and OUTLAST regulator constructs were enumerated using an automated cell counter. T cell suspensions were adjusted to 1 x 10^4 Ror1 CAR+ T cells/100 uL in R10 media (based off of post-bead purification flow staining) and added directly to H1975-mKate2 pre-seeded wells to initiate round 1 of tumor challenge at an Effector:Target (E:T) ratio of 1:2 (Ror1 CAR+ T cells:H1975-mKate2). Plates were then placed in an IncuCyte S3 Live-Cell Analysis System (Sartorius) housed within a humidified incubator (37℃, 5% CO_2_), and images were acquired every 6 hours in red fluorescence and phase-contrast channels. Sequential tumor re-challenges were performed every 3 to 4 days for a total of 8 rounds. On the morning of each re-challenge, fresh H1975-mKate2 target seeded plates were prepared at 2 x 10^4 cells/well in 100uL R10 media and incubated for 2–3 hours to permit cell adherence. Co-culture cell suspensions from the preceding challenge round were gently resuspended by pipetting and transferred to 96-well V-bottom plates. Plates were centrifuged at 300 x g for 10 minutes at room temperature, and supernatants were decanted. Cell pellets were resuspended in 150 uL of fresh R10 media per well, then 100 uL were transferred into the newly prepared target cell seeded plates. Plates were immediately returned to the IncuCyte system to resume real-time imaging. Target cell killing was quantified continuously over time by tracking total mKate2+ (red fluorescence) cell counts, with values normalized independently to baseline (t = 0 hours) at the start of each re-challenge round.

##### *In vitro* live-cell chronic stimulation and subsequent 2D cytotoxicity assay

To induce chronic antigen stimulation prior to functional evaluation, target H1975-mKate2 cells were seeded into 6-well tissue culture-treated plates at 1 x 10^6 cells/well in 1.5 mL of R10 media and incubated for 2–3 hours at 37℃, 5% CO_2_ to allow cell adherence. Freshly prepared Ror1 CAR-T cells were enumerated using an automated cell counter. For single-vector (all-in-one) transduced T cells, suspensions were adjusted to 1 x 10^6 ROR1 CAR+ T cells/1.5 mL of R10 media. For dual-transduced and bead-purified T cells (which exhibited >90%ROR1 CAR+ purity), T cells were adjusted to 1 x 10^6 total viable T cells/1.5 mL of R10 media. 1.5 mL of adjusted T cell suspensions were then seeded into prepared H1975-mKate2 plates at an effector-to-target (E:T) ratio of 1 (3 mL/well), then subjected to chronic stimulation with live H1975-mKate2 target cells over a 7-day period. On days 2 and 4 post-co-culture setup, co-cultures were reset as follows: primary T cells were harvested, counted, and re-plated onto freshly prepared plates containing 1 x 10^6 live H1975-mKate2 cells/well to re-establish a 1:1 E:T ratio. Following 7 days of continuous chronic stimulation, persistent T cells were harvested, counted for viability, and transferred to secondary cytotoxicity assays against fresh 2D H1975-mKate2 target cells. Target cells were pre-seeded in 96-well flat-bottom plates at 2 x 10^4 cells/well in 100 uL R10 media. Chronically stimulated viable T cells were added at graded E:T ratios of 1:2, 1:4, 1:8, and 1:16 (Viable T cells: H1975-mKate2). Microplates were placed in an IncuCyte S3 Live-Cell Analysis System (Sartorius) housed within a humidified incubator (37℃, 5% CO_2_), and images were acquired every 6 hours for 5–7 days in red fluorescence and phase-contrast channels. Target cell killing was quantified over time by tracking the total mKate2+ (red fluorescence) cell count, normalized to baseline (t = 0). To assess effector cytokine release, 50 uL/well of supernatants were harvested 18–24 hours post-killing assay setup and stored at −80℃ for subsequent quantification of secreted IFN*_γ_* and IL-2 by Meso Scale Discovery (MSD) electrochemiluminescence immunoassay.

##### *In vitro* chronic stimulation on immobilized recombinant antigen and subsequent 3D tumor spheroid assay

To evaluate T cell functionality following sustained, antigen-specific TCR engagement in the presence of adhesion ligands, engineered primary human T cells expressing a PRAME-specific TCR (Vb8+/Dextramer+) were subjected to chronic stimulation using immobilized recombinant protein complexes. At least one day prior to chronic stimulation setup, 24 well plates were coated with immobilized recombinant protein matrices. In order to capture biotinylated PRAME peptide-MHC (pMHC) complexes, 24 well tissue culture plates were initially coated with 300 uL/well NeutrAvidin (5 ug/mL in PBS), then washed twice with 300uL PBS. After washing, 300 uL/well of a biotinylated PRAME pMHC (20nM), recombinant human CD58 (2 ug/mL), and recombinant human ICAM-1(2 ug/mL) solution diluted in PBS was added. Antigen coated plates were wrapped in parafilm and stored at 4℃ for at least 18-24 hours until ready to use for chronic stimulation. The day before chronic stimulation setup, cryopreserved T cells were thawed and rested overnight (16-24 hours) at 37℃, 5% CO_2_ in T cell media supplemented with human IL-2, IL-7, and IL-15. The following day rested T cells were harvested, enumerated, and normalized for TCR expression based on PRAME-specific dextramer positivity (Dextramer+) determined by flow cytometry pre-cryopreservation. Coating solutions were aspirated from the 24-well plates, and normalized Dextramer+ T cells were seeded into antigen-coated wells in cytokine-supplemented T cell media to initiate chronic stimulation. Cultures were maintained at 37℃, CO_2_ for a total of 6 days. On Day 3, chronic co-cultures were reset by gently resuspending and harvesting the total cell suspension from each well without cell-number normalization. Cells were pelleted by centrifugation at 300 x g for 10 min, resuspended in fresh cytokine-supplemented T cell media, and transferred onto freshly prepared NeutrAvidin:biotin-pMHC/CD58/ICAM-1 matrix plates. To prepare 3D tumor targets for downstream cytotoxicity assays, H1650-mKate2 cells were seeded on Day 3 into PrimeSurface 96U 3D Ultra-Low Attachment Cell Culture Plates (S-Bio) at 1 x 10^4 cells/well (100 uL/well). Microplates were centrifuged at 1,000 x g for 10 min to promote cell compaction and incubated for 3 days at 37℃, CO_2_ to allow tight spheroid assembly. Following 6 days of continuous chronic stimulation, an aliquot of persistent T cells were harvested and flow staining was performed with a panel targeting T cell lineage, differentiation, and exhaustion markers (including anti-CD45RA-BUV395, anti-CD4-BUV496, anti-CD62L-BUV615, anti-CD27-BUV737, anti-CD8-BV421, anti-CD39-BV510, anti-CD28-BV711, anti-LAG3-Super Bright 780, anti-Vb8-APC, anti-TIGIT-PE, anti-PD-1-PE-Cy7, Fixable live/dead eFluor780, Brilliant Stain Buffer Plus). Reported datasets only include TIGIT and PD-1 as indicated in **Figure 6**. The remaining T cells were counted using an automated cell counter and diluted to normalized CD8+Vb8+ T cell concentrations. CD8+Vb8+ normalized T cells were then transferred to secondary cytotoxicity assays against prepared H1650-mKate2 spheroids at graded Effector-to-Target (E:T) ratios of 1:5 and 1:10 (CD8+Vb8+ T cells:H1650-mKate2). Microplates were placed in an IncuCyte S3 Live-Cell Analysis System (Sartorius) housed within a humidified incubator (37℃, 5% CO_2_), and images were acquired every 6 hours for 5–7 days in red fluorescence and phase-contrast channels. Target cell killing was quantified over time by tracking the Integrated Red Fluorescence Intensity/well (RCU x um^2/well) normalized to baseline (t = 0). To assess effector cytokine release, 50 uL/well of supernatants were harvested 18–24 hours post-killing assay setup and stored at −80℃ for subsequent quantification of secreted IFN*_γ_* and IL-2 by Meso Scale Discovery (MSD) electrochemiluminescence immunoassay.

##### Quantification of secreted IFN-*_γ_* and IL-2 by Meso Scale Discovery (MSD)

Concentrations of secreted human IFN*_γ_* and IL-2 were quantified using a custom multiplex electrochemiluminescence immunoassay (Meso Scale Discovery) according to the manufacturer’s protocols. Plates were read on an MSD automated plate reader controlled by Methodical Minds software. Absolute cytokine concentrations (pg/mL) were interpolated from 4-fold, 8-point calibrator standard curves using MSD Discovery Workbench analysis software.

##### *In vivo* experiments

*In vivo* studies were carried out by Outpace Bio staff at a CRL Accelerator and Development Lab (CRADL) in Seattle, WA using either NOD.Cg-*Prkdc^scid^Il2rg^tm1Wjl^*/SzJ (NSG) mice (Jackson Laboratory) or NOD-*Prkdc^em26Cd52^Il2rg^em26Cd^*^22^/NjuCrl (NCG) mice (Charles River Laboratories).

The H1975 ROR1 CAR-T tumor efficacy study was carried out using NSG mice bearing tumors implanted subcutaneously at 5×10^6^ cell/mouse. The H1975 tumor cells were resuspended in 100uL of PBS and mixed 1:1 with 100uL of Matrigel (Corning) for a 200uL injection in the right flank. After implant, mice were monitored twice weekly for body weight and tumor volume performed using caliper measurements and the formula (width^2^ x length)/2 was used for calculating tumor volume. Nine days post implant, mice were randomized into study groups of 5 with mean volumes falling between 98mm^3^-102mm^3^. Frozen vials of engineered CAR cells from two different human donors were thawed and formulated to contain either 1×10^6^ or 2×10^6^ total CAR^+^ cells and injected intravenously via tail vein in 200uL of saline (Cytiva). Mock cells were dosed to match the maximum total cell number of the highest dose group for each donor. Animals exited study when either a maximum tumor burden of 2000mm^3^ was reached or when body weight dropped below 80% of baseline, whichever occurred first. Peripheral blood collections occurred via the submental route (100uL) into EDTA coated tubes (BD) and processed for flow cytometry. For staining, 50uL of whole blood was mixed with 50uL of staining antibody cocktail for 30 minutes at 4°C then incubated for 15 minutes with 1.6mL of 1X RBC lysis/fixation solution (BioLegend). Samples were washed with Cell Staining Buffer (BioLegend) and prepared in 150uL for analysis on a ZE5 cytometer (BioRad) and analyzed using FlowJo software (TreeStar, Inc; v10). Blood samples were stained using hCD45-APC (Clone 2D1, BioLegend), CD3-BV650 (Clone OKT3, BioLegend), CD4-BUV496 (Clone SK3, BD), CD8-BV785 (Clone RPA-T8, BioLegend), CD39-BV510 (Clone A1, BioLegend), EGFR-BV421 (Clone AY13, BioLegend), CD45RA-BUV395 (Clone HI100, BD), CD62L-BUV615 (Clone DREG-56, Thermo Scientific), TIGIT-PE (Clone A15153G, BioLegend), PD1-PE-Cy7 (Clone J105, Thermo Scientific) and Brilliant Stain Buffer^+^ (BD). Precision Count Beads (BioLegend) were included at known concentration for absolute quantification of cells.

The H226-A*02:01 PRAME^+^ tumor efficacy study was carried out using NCG mice bearing tumors implanted subcutaneously and monitored as previously described. On day 25 post tumor implant mice were randomized into study groups of 5 with an average tumor volume of 147mm^3^.

##### Flow cytometric phenotyping for metabolic/mitochondrial Profiling

Rested dual-transduced and bead-purified T cells were harvested into a 96-well U-bottom plate and washed twice with PBS (500 x g, 2 min). Cells were resuspended in 50 uL of PBS alone, or 50 uL of PBS containing carbonyl cyanide m-chlorophenyl hydrazone (CCCP; 100 uM final concentration) as a depolarization control for mitochondrial membrane potential (Δ*_Ψ_*_m_). Plates were incubated for 30 min at 37℃, 5% CO_2_. Without intermediate washing, 50 uL of a 2x dye cocktail containing 2-NBDG (100 uM final concentration), MitoTracker Deep Red FM (MTDR; 1:1000 final dilution), and Tetramethylrhodamine Methyl Ester (TMRM; 100 nM final concentration) in PBS was added directly to each sample (final staining volume 100 uL). Cells were incubated at 37℃, 5% CO_2_ for an additional 45 min. Following metabolic and mitochondrial dye staining, cells were washed twice with Cell Staining Buffer (CSB; BioLegend) then stained with a fluorophore-conjugated surface antibody and viability dye cocktail (anti-CD8-BUV395, anti-CD4-BUV496, anti-EGFR-BV711, anti-CD19-FITC, Fixable Viability Dye eFluor 780, and Brilliant Stain Buffer Plus) for 20-30 min at room temperature in the dark. Cells were subsequently washed twice with CSB and resuspended in CSB for acquisition. Flow cytometric acquisition was performed on a ZE5 Cell Analyzer (Bio-Rad). All flow cytometry data were analyzed using FlowJo software (BD Biosciences, v10).

Data are reported as mean fluorescence intensity (MFI) normalized to the trimer-only negative control within live EGFR+ gated cells as follows:

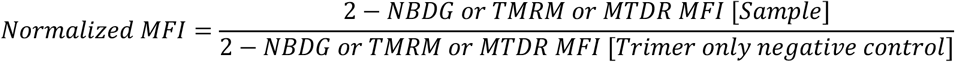

Data represent exploratory profiling performed on n = 1 healthy human donor.

### QUANTIFICATION AND STATISTICAL ANALYSIS

#### *In vivo* experiment statistical analysis

Statistical analyses were GraphPad Prism (v10.2.2). For in vitro and in vivo killing assays, tumor burden was summarized by calculating the area under the curve (AUC). For studies with three or more donors, a linear mixed-effects model was used to compare treatment groups, with donor as a random effect. Survival curves were analyzed using the log-rank (Mantel-Cox) test.

